# A *De Novo* Algorithm for Allele Reconstruction from Oxford Nanopore Amplicon Reads, with Application to *CYP2D6*

**DOI:** 10.1101/2025.11.06.687066

**Authors:** Scott D. Brown, Lisa Dreolini, Agata Minor, Michelle Mozel, Nancy Wong, Sharon Mar, Amanda Lieu, Maimun Khan, Amanda Carlson, Monica Hrynchak, Robert A. Holt, Perseus I. Missirlis

## Abstract

The Oxford Nanopore Technologies’ sequencing platform offers a path towards bedside genomics, producing long reads that can completely cover a gene of interest, and thus detect any known or novel variant the gene contains. However, the analysis of these long reads to identify actionable genotypes remains challenging and typically requires customization depending on the target gene. Here, we describe a generic algorithm to accurately reconstruct allele sequences derived from long-reads of genomic-amplicon origin. Rather than calling variants directly from these long-reads, our method takes a “sequence-first” approach, performing an unbiased reconstruction of the underlying amplicon sequences to generate high-confidence reconstructed allele sequences. This is done without user input of the expected target gene, allowing for any source amplicon to be reconstructed. These high-confidence reconstructed allele sequences are then compared to the genomic reference sequence of the gene to infer the specific diplotype present in the sample. This approach is agnostic towards the number of genes and alleles present and readily detects novel variants. We demonstrate our approach using three independent data sets for *CYP2D6*, a diverse and complex gene with over 175 known alleles of clinical significance affecting drug dosing. We show how our approach can accurately recover validated *CYP2D6* diplotypes from 20 Coriell samples sequenced using different primer sets, on different Oxford Nanopore Technologies flow cell versions, and to different depths. This includes inferring occurrences of copy number variation from relative abundances of each allele, a critical factor for ascribing functional effects to a diplotype. Further, we demonstrate our approach’s utility for other genomic regions, including *HLA*.

## Background

Gene diversity can significantly impact the therapeutic effect of drugs in patients. One such example is the cytochrome P450 (CYP) superfamily of enzymes that play a key role in drug metabolism and can result in either supratherapeutic or subtherapeutic serum drug concentrations (McDonnell and Dang, 2013). Genotyping of genes prior to treatment can inform drug selection, dosing decisions, and avoidance of adverse side effects (Chang *et al*., 2015). *CYP2D6* is one of the most clinically significant genes and *CYP2D6* genotype-guided dosing exists for a number of indications, including anti-fungal treatment in hematopoietic stem cell transplantation, breast cancer chemotherapeutics, tricyclic antidepressants, and opioids for pain management (Teusink *et al*., 2016; Kiyotani *et al*., 2008; Hicks *et al*., 2017; Somogyi *et al*., 2015). As such, a method for obtaining a fast and accurate readout of a patient’s *CYP2D6* genotype is of great importance. *CYP2D6* is highly polymorphic; currently there are over 175 known functional alleles (each designated as a “star-allele”, with *CYP2D6*1* being the reference), many of which differ from another allele by only a single base substitution (Whirl-Carrillo *et al*., 2021, 2012), and new alleles are constantly being added as they are discovered.

Traditional methods to interrogate genotypes are generally slow and expensive; typically relying on custom sets of PCR reactions to interrogate specific sites along the length of each gene of interest. More contemporaneous approaches have leveraged high-throughput sequencing data, either whole genome (Chen *et al*., 2021; Deserranno *et al*., 2023), or amplicon-based (Hari *et al*., 2023; Liau, Cree, *et al*., 2019; Liau, Maggo, *et al*., 2019; Qiao *et al*., 2016; Ammar *et al*., 2015; Charnaud *et al*., 2022). These methods often make use of Pacific Biosciences or Oxford Nanopore Technologies (ONT) long reads, allowing the entire gene to be sequenced in a single read, and thus, facilitating accurate DNA sequencing and haplotyping with a single assay. Regardless of sequencing strategy, these methods take a “variant-first” analysis approach, relying on variant calling from the sequence reads first, followed by phasing of the variants into haplotypes.

We sought to develop a method of diplotyping genes that uses a “sequence-first” rather than “variant-first” approach. We aimed to perform allele sequence reconstruction from long read sequences by generating high-confidence consensus sequences of each allele present in the original sample, then genotyping each resulting high-confidence consensus sequence to assign a haplotype. Further, we ensured our method was unbiased and generalizable, not being restricted to a specific gene. As such, our only input requirements are that the dataset is generated from genomic amplicon sequencing using ONT long reads and the dataset contains reads which completely cover the genomic regions of interest. No information is needed *a priori* regarding the specific genomic target, nor the number of targets. We demonstrate the utility of our approach using ONT sequencing data from *CYP2D6* amplicons and scripts that automate *CYP2D6* haplotyping of reconstructed allele sequences to call the most likely star-allele and identify any discrepant, potentially novel, variants.

## Results

### Selection of CYP2D6 amplicon

The *CYP2D6* gene locus is 4.3 kb long and is located on the negative strand of chromosome 22 (chr22:42,126,499-42,130,810; hg38). Variants affecting CYP2D6 enzyme function may fall anywhere within this region, with the range of currently known variants (insertions, deletions, and single nucleotide variants) spanning chr22:42,126,578-42,130,783 (Whirl-Carrillo *et al*., 2021, 2012). We used primers previously developed by Ammar et al. (Ammar *et al*., 2015) to selectively amplify a 5.1 kb region of *CYP2D6* (chr22:42,126,072-42,131,142), and not the nearly identical *CYP2D7* pseudogene (**Figure 1**).

**Figure 1.**
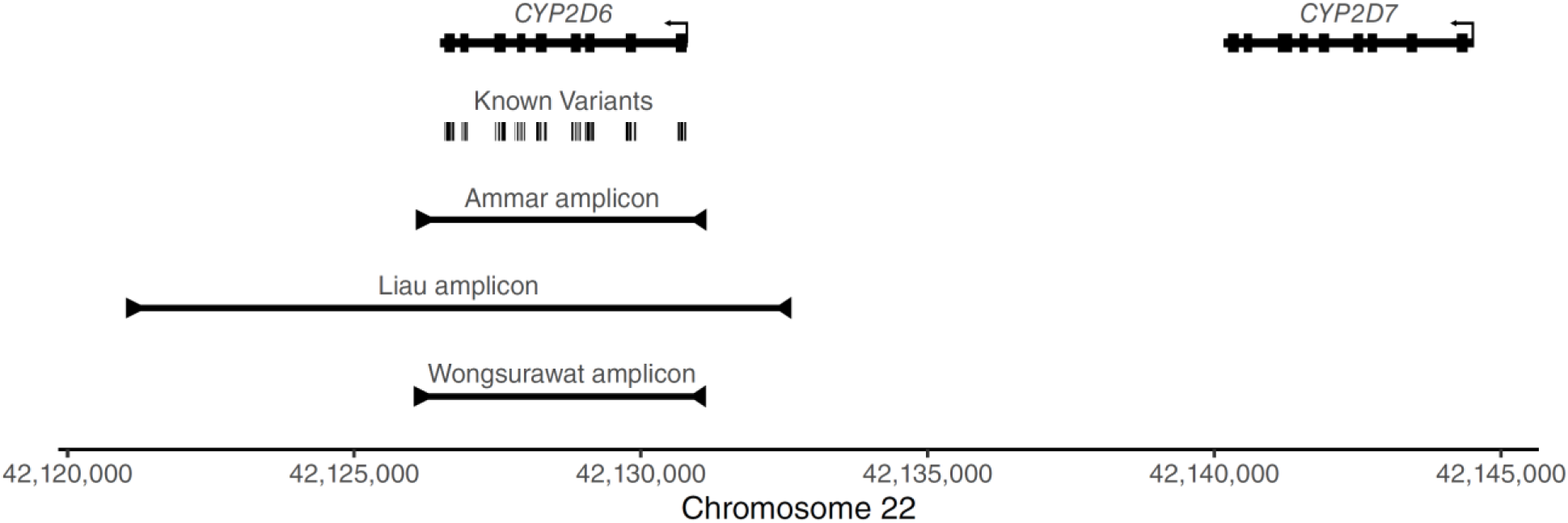
Graphical summary of the genomic region encompassing *CYP2D6*. The *CYP2D6* gene is shown, with larger boxes for the exons. The *CYP2D7* pseudogene is shown upstream of the *CYP2D6* transcriptional start site. All known functional variants used by ClinPGx (Whirl-Carrillo *et al*., 2021, 2012) to define specific *CYP2D6* alleles are shown as ticks. The three amplicon regions assessed in our work are shown as individual tracks.

### Oxford Nanopore Technologies sequence generation

Genomic DNA from three Coriell samples with known *CYP2D6* haplotypes (NA02016; *\*2xN/*17*, NA10005; *\*17/*29*, NA17300; *\*1/*6*) were used. Sequencing libraries were constructed using the ONT Ligation Sequencing Kit V14. We loaded each amplicon library onto a separate R10.4.1 Flongle flow cell, each being run for 24 h on a MinION Mk1b USB sequencer.

Base calling of the raw .fast5 files was performed using the GPU-enabled version of Guppy (a base caller provided by ONT). No quality filter was used at this stage, as quality filtering was performed at the per-base level later in our analysis pipeline. Omitting the quality filter allows for high-quality bases in lower-quality reads to still contribute information, maximizing the depth of coverage at all positions. Porechop was used to remove any adapter sequences and split chimeric reads. We obtained 87,906, 56,948, and 50,389 total reads from NA17300, NA10005, and NA02016 respectively. The average read length was 1,736 bases (± 2,078.35) (**Figure 2**). We observed a median 8.5 % error rate in these reads (range 7.0 – 9.0 %).

**Figure 2.**
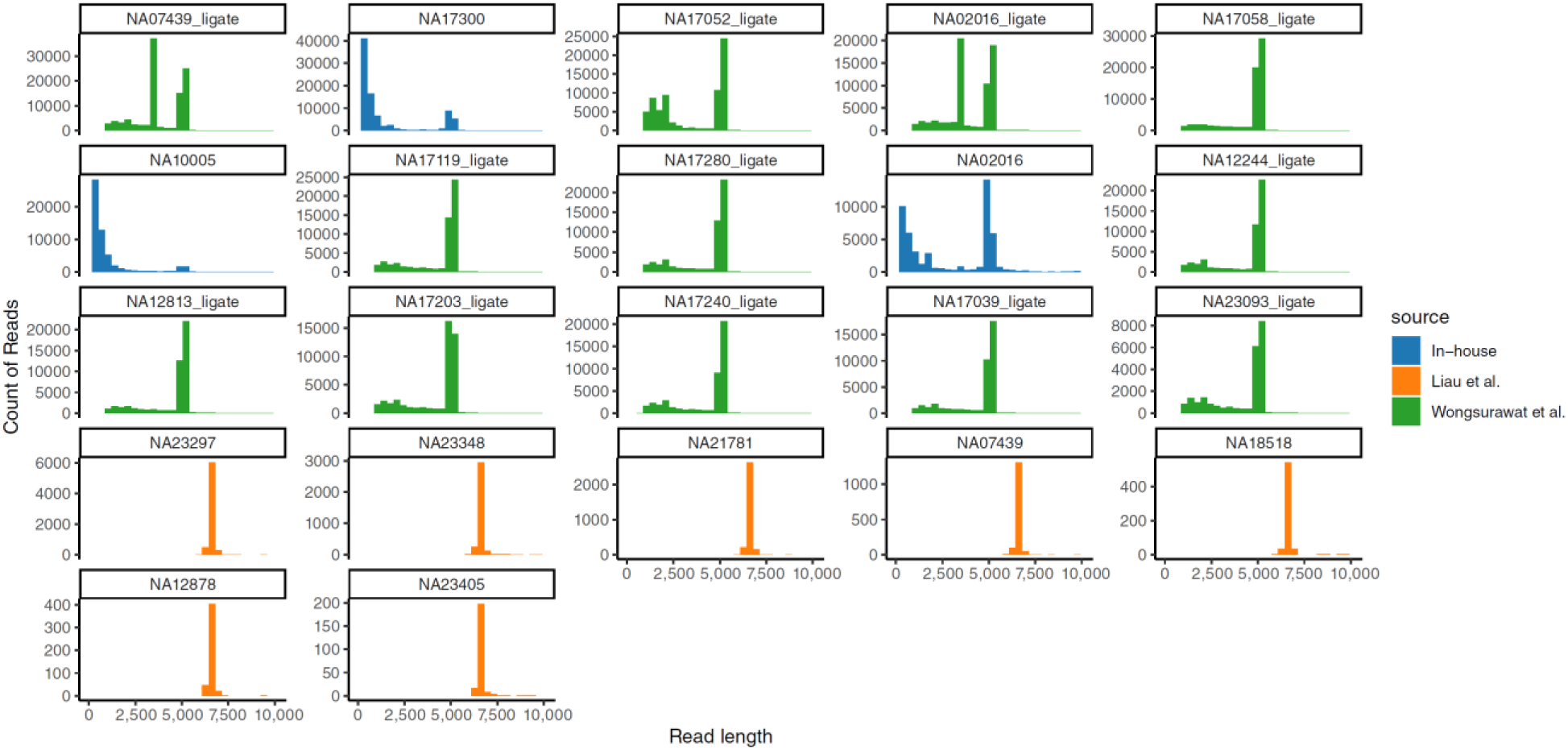
Distribution of read lengths for all samples. Samples ordered by decreasing numbers of total reads. In all cases, a population of reads at the expected 5.1 – 6.6 kb range is seen. In-house generated samples (blue) show higher than expected numbers of short reads, suggestive of degraded DNA amplicons during library preparation. The x-axis was truncated at 10,000 bp, removing an average of 0.37 % of reads per sample from display.

### Inclusion of existing *CYP2D6* sequence data sets to test different flow cells, sequence qualities, and amplicons

To validate our approach on different ONT flow cell versions, *CYP2D6* amplicons, sequencing depths, and haplotypes, we obtained data from Liau et al. (Liau, Maggo, *et al*., 2019) for seven additional Coriell samples. Liau et al. generated a 6.6 kb amplicon for *CYP2D6* **(Figure 1)**, which covered more of the upstream and downstream regions of *CYP2D6,* and sequenced these on older MinION R9.4 flow cells, yielding an average of 2,292 reads per sample, and having a median error rate of 12.1 % (range 11.9 – 12.5 %). This larger 6.6 kb amplicon should have a greater ability to detect gene duplication events, being more likely to span the junction site of a duplicated copy, though the exact location of this junction is unknown. Of note, Liau et al. also sequenced an even larger 8.6 kb amplicon for samples with confirmed duplications, covering the repetitive region downstream of *CYP2D6* (Gaedigk *et al*., 2019), but this sequence data was unavailable.

Data for twelve additional Coriell samples were obtained from Wongsurawat et al. (Wongsurawat *et al*., 2025), with an average of 55,068 reads per sample, and a median error rate of 5.1 % (range 4.7 – 5.7 %). Wongsurawat et al. generated a 5.1 kb amplicon using primers from Puaprasert et al. (Puaprasert *et al*., 2018), covering a region similar to Ammar et al., and were multiplexed on MinION R9.4.1 flow cells. Wongsurawat et al. included two additional primer sets, one designed to identify gene deletions by producing a 3.5 kb amplicon only upon gene deletion (otherwise priming at sites 15 kb apart; too large to be efficiently amplified), and another designed to amplify a 3.5 kb region between duplicated copies of *CYP2D6* (with priming sites on either side of *CYP2D6* facing “outwards”). This data was generated using two different library construction kits; the Rapid Barcoding Kit and the Ligation Sequencing Kit. We only used Ligation Sequencing Kit data for our analysis. Combined with our in-house generated data, we have compiled 22 ONT sequence datasets covering 20 distinct Coriell samples.

### Assigning reads to inferred amplicons based on whole genome alignment

We designed our method to perform an unbiased reconstruction of allele sequences. We base this on an assumption that every amplicon present in the sequence library will have a subset of reads supporting that amplicon. As such, rather than supplying the algorithm with the anticipated amplicon, we used the sequence reads to infer the genomic loci that were amplified. To achieve this, we aligned all reads to the human reference genome (hg38) using minimap2, allowing for only the single best mapping position of each read. Supplementary alignments, or split reads, were included in order to detect large deletions or structural variants. The depth of coverage across the whole genome was assessed, and the regions with high coverage were taken to be representative of amplicons present in the source sequencing library (**Figure 3A**). Importantly, this approach does not assume any predefined number of amplicons, allowing this approach to work, theoretically, for multiplexed samples of any number of amplicons. All of our samples yielded an inferred amplicon region at the expected *CYP2D6* locus, and six samples yielded at least one second, distinct inferred amplicon region (**Supplementary Table S1**). Two of these non-*CYP2D6* inferred amplicons (NA02016_ligate, NA07439_ligate) were from the region between *CYP2D6* and *CYP2D7*, corresponding to the duplication-detecting amplicon from Wongsurawat et al. One (NA17203_ligate) was from the *CYP2D7* locus, possibly derived from a *CYP2D6*-*CYP2D7* hybrid allele, where the majority of the amplicon sequence was derived from *CYP2D7*. Three (NA02016, NA10005, NA17052_ligate) were from elsewhere in the genome. NA02016 and NA10005 produced short (mean 609 bp) off-target amplicon regions from chromosomes 4, 9, 12, and 20. NA17052_ligate had an off-target amplicon fromfrom the *SORCS2* locus on chromosome 4. Careful inspection of the identified *SORCS2* locus sequence shows it contains a string of 15 contiguous bases at the start (GCCCTGGGAGGTAGG), and 13 contiguous bases at the end (GAAGCCTGCAAAG), which exactly match sequence within the *CYP2D6* locus, and are present in the primers used by Wongsurawat et al. All inferred amplicons are carried forward in the analysis as possible loci of interest.

**Figure 3.**
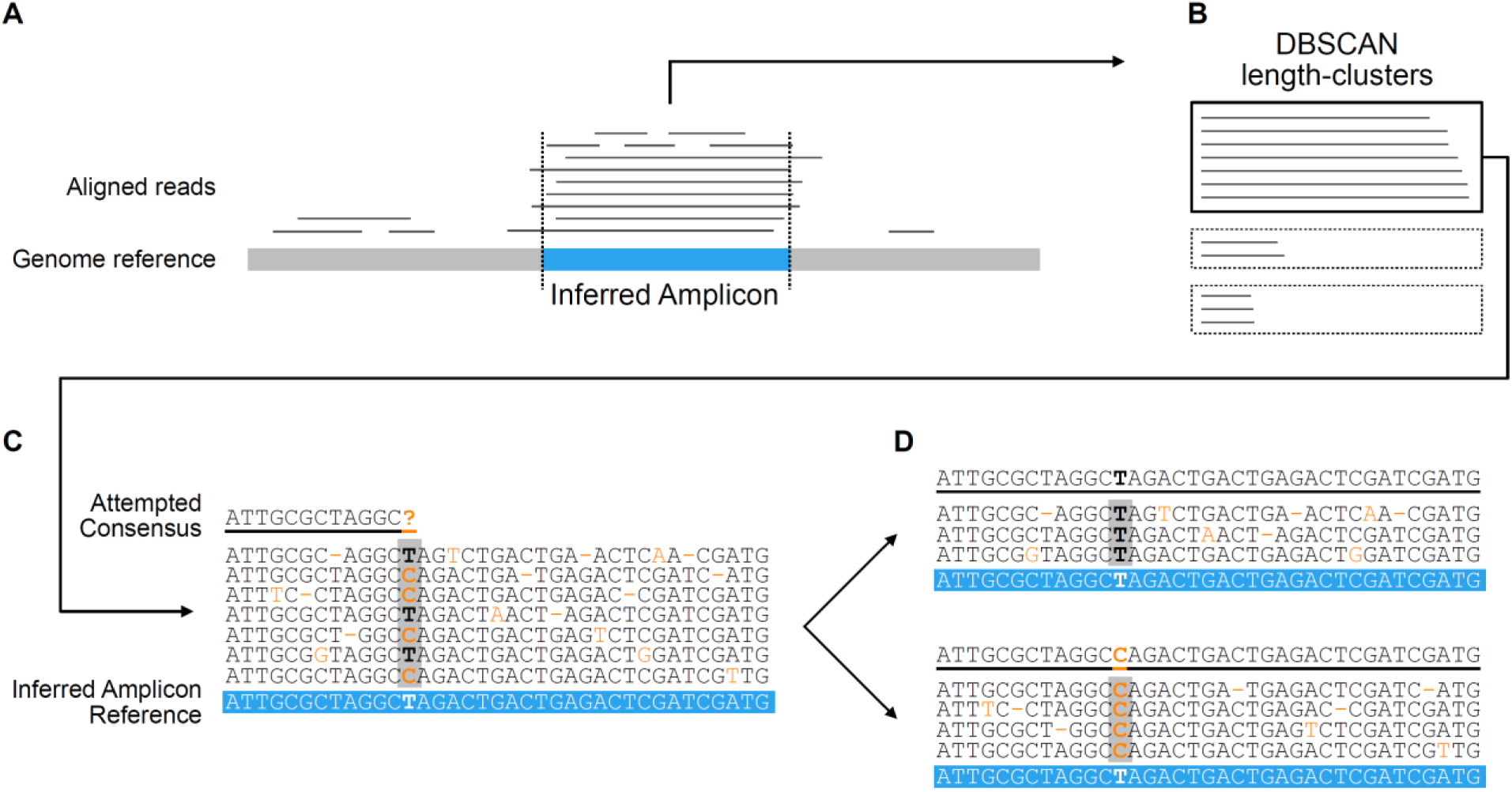
Schematic overview of algorithm. (A) Sequence reads are aligned to the genome, and regions of high depth are extracted and inferred as the source amplicon (blue). (B) These reads are clustered by length using the DBSCAN algorithm, and the most abundant cluster(s) are selected for allele reconstruction (solid outlined box). (C) Selected reads are aligned to the inferred amplicon reference sequence (extracted from the genomic reference; blue), and each position of the alignment is assessed sequentially, building a consensus sequence base by base (top underlined sequence). Variants as compared to the reference are coloured orange. If a position is encountered as appearing heterozygous (having roughly equal support for each of two or more bases), resulting in an unknown consensus base, the reads are partitioned into subsets based on the base they support at that position (D). The reads in each subset are used, independently, to re-attempt allele reconstruction. In the example shown, this yields two consensus sequences (top underlined sequence), each with homozygous support for each base. Small indels are treated the same way as bases, with partitioning occurring similarly for reads supporting and not supporting the indel.

### Reads clustered into groups based on length

In our in-house generated data, we observed a higher than expected frequency of short reads (**Figure 2).** On average, 60.8 % of reads were under 1 kb, compared to 0 % for data from Liau et al. (only aligned reads were provided) and 0.08 % from Wongsurawat et al. We aligned our short reads to the *CYP2D6* reference sequence, demonstrating that they mapped to random positions spread across the amplicon (**Supplementary Figure S1**). This suggested the short reads were derived from fragmented amplicons during library construction, and were not representative of *bona fide* short amplicons within the sample. Liau et al. describe seeing many short reads in their first run, which prompted them to use modified beads to remove any DNA fragments shorter than 3-4 kb (Liau, Maggo, *et al*., 2019), which could be an approach we take in the future. We assert that full-length (non-fragmented) reads which were derived from the same amplicon would all have the same length (or highly similar lengths when allowing for sequence errors). To select the subset of reads for each inferred amplicon most likely to be informative, we performed length-clustering on the reads from each inferred amplicon using the DBSCAN algorithm (Schubert *et al*., 2017) (**Figure 3B**). We performed a parameter space exploration of the two DBSCAN parameters, EPS and Minimum Neighbours (see Materials and Methods), which revealed an inverse relationship between the variability of read lengths and size of the resulting clusters. Low EPS and high minimum neighbours result in a more strictly defined cluster, ensuring all reads have a similar length, at the sacrifice of number of reads, whereas high EPS and low minimum neighbours allow larger clusters with more variable lengths. Since the purpose of length-clustering here is to identify subsets of reads from the same amplicon, we chose the stricter parameter values to avoid using shorter, degraded reads. After clustering, we first selected the cluster that had reads with a mean length that was closest to the source inferred amplicon, regardless of abundance, representing reads that fully covered the amplicon. Secondly, we also assessed any highly abundant clusters, based on the premise that there may be other smaller or larger amplicons that would otherwise go undetected (**Supplementary Table S1**). All selected clusters were used for subsequent allele sequence reconstruction.

### Allele sequence reconstruction performed through unbiased, recursive consensus sequence generation

At this stage, we had obtained subsets of length-matched reads which were likely to have been derived from the same locus. These subsets, however, contained a mix of reads supporting each of the underlying haplotypes present in the original sample. If all sequence reads were error-free, this would have been a trivial exercise to partition reads based on their exact sequence. Due to sequencing errors, reads derived from the same haplotype did not share the exact same sequence. In order to reconstruct the underlying allele sequences, we took an empirical approach for determining which allele sequences the reads supported. This was achieved by finding partitions of the reads such that all reads in a partition supported the same sequence (while allowing for some noise due to sequence errors). First, we attempted to build a consensus sequence from all of the reads in the set, using similar logic to our previous work (Brown *et al*., 2023). Briefly, the reads were aligned to the inferred amplicon reference sequence, and the pileup of base support at each position of the alignment was assessed, sequentially, to call a consensus base at that position. If a position was reached that did not have sufficient evidence of a homozygous base (homozygosity defined as the top two bases having a ratio of at least 2:1; see Materials and Methods) (**Figure 3C**), the reads were partitioned into subsets defined by the observed bases at this heterozygous position (**Figure 3D**). The consensus sequence generation was restarted from the beginning, recursively, on each of these resulting subsets. This process continued until a homozygous consensus sequence was successfully generated from a subset of reads, or the size of a subset fell below the lower limit threshold for reporting (5% of the total starting read set of length-clustered reads, selected empirically).

Our approach considered base quality in two ways. In order for a base to contribute to the consensus sequence, the base must have had a minimum quality of 10 (selected empirically). If this criterion was not met, a secondary criterion was tested, requiring a quality score above the 33^rd^ percentile of observed qualities at that position (selected empirically). This ensured that positions with lower-quality bases across all reads were not completely lost from the consensus due to lack of coverage, while still ensuring that the consensus sequence was built using the best quality bases available at each position. Strand-specific errors are also common in ONT data (Krishnakumar *et al*., 2018; Branton *et al*., 2008), resulting in a specific sequencing error occurring primarily on one strand. When assessing all reads without considering the strand, such errors could appear as a heterozygous position. To avoid erroneously calling a position heterozygous, we assess, separately, the positive and negative strand reads at each position, requiring support of heterozygosity from both strands independently in order to classify a position as truly heterozygous. All reconstructed consensus sequences are available in **Supplementary File S1**.

Focusing on the results for read sets derived from the correct genomic locus (*CYP2D6*) and of the correct mean length (5.1 - 6.6 kb) for the 22 samples, all yielded at least two distinct consensus sequences (range 2-27). While having two sequences is expected for a diploid genome, having greater than two consensus sequences could support gene duplication events where the copies have sequence differences. This could also, however, be from a common sequencing error that is abundant enough to result in a consensus sequence. For samples with a large number of consensus sequences, the distribution of read depths was highly skewed, typically with only 2 consensus sequences having the majority of the reads. This is consistent with these lower-abundance consensus sequences arising from common sequencing errors and not *bona fide* amplicons. Relative read depths may help distinguish these two cases, as described later.

All three samples sequenced by our group had length-clusters that were shorter than the expected amplicon size, but were abundant enough to have allele sequence reconstruction attempted (**Supplementary Table S1**). This is likely an artefact from our high rate of short reads. One yielded two consensus sequences mapping to exon 9 and downstream of *CYP2D6* (NA02016), one yielded two consensus sequences mapping to exons 8-9 of *CYP2D6* (NA10005), and one yielded a consensus sequence mapping to exons 4-6 of *CYP2D6* (NA17300). This supports the notion that these short reads were derived from degraded DNA template occurring during library construction. Indeed, when inspecting these short consensus sequences, they match the respective regions of the long consensus sequences from the same sample (**Supplementary Figure S2**). Two samples had allele sequence reconstruction attempted for reads that did not originate from the *CYP2D6* locus or supplementary amplicons. As described above, these are most likely derived from off-target amplification from the genome during PCR, but could also be the result of *CYP2D6*-*CYP2D7* hybrid alleles (in the case of the *CYP2D7*-derived consensus sequence in NA17203_ligate). The *SORCS2*-derived consensus sequences were run through BLASTn (Sayers *et al*., 2025), and both mapped back to the *SORCS2* locus, supporting that these reads were from off-target amplification.

### Confirmation of deletion detection through simulated data

To confirm our allele sequence reconstruction would generate a consensus sequence supporting a large deletion relative to the genomic reference, should one exist, we simulated long reads representing our amplicon sequence with the entire *CYP2D6* sequence deleted (see Materials and Methods). Encouragingly, our pipeline output a consensus sequence that perfectly matched the source template sequence used to simulate the reads, despite the reads containing a 4.3 kb deletion relative to the human reference genome. This demonstrates that our algorithm will accurately produce a consensus sequence representative of the sequence the reads support. ***Automated reporting of the most likely* CYP2D6 *star alleles and putative novel variants*.**

To identify the specific *CYP2D6* alleles that each resulting consensus sequence supported, we first aligned each consensus sequence to the *CYP2D6*1* reference sequence. All variants (indels and mismatches) were enumerated, and those occurring within coding regions or splice sites were used for *CYP2D6* allele determination. We automated this process by using the *CYP2D6* Allele Definition Table from ClinPGx (Whirl-Carrillo *et al*., 2021, 2012). Briefly, for each known star allele, all canonical variants were parsed from the ClinPGx definition table. For alleles with variants that have multiple possible bases allowed, some of which may include the reference wildtype base, all possible allowed bases are stored. Then, the set of identified variants for the consensus sequence was compared to the set of all required or possible variants from all known alleles, and each known star allele was given a score by summing the number of disagreements in the identified variants from the consensus sequence. Thus, a consensus sequence that had exactly the same variants as a defined star-allele would receive a score of zero for that allele. Silent mutations found in the consensus would negatively impact the score across all alleles equally. The allele with the lowest score was returned as the best candidate allele assignment for the consensus sequence. In the event of an imperfect match, missing and/or extra variants in the consensus sequence compared to the candidate star allele reference were also returned, allowing for manual review of the candidate allele assignment (**Table 1**). In this way, novel variants and alleles can also be readily identified, rather than simply the best match to known alleles.

**Table 1.**
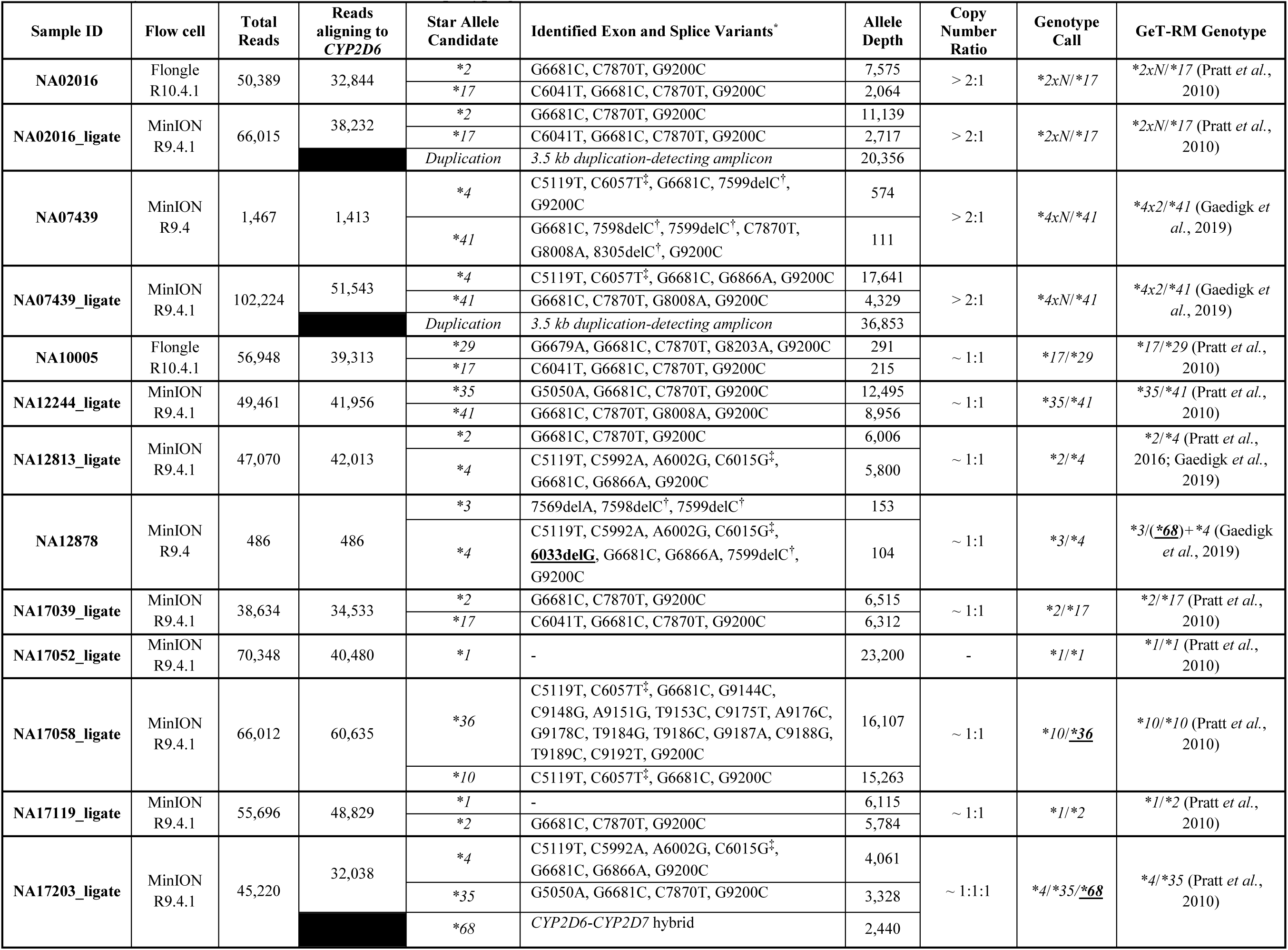
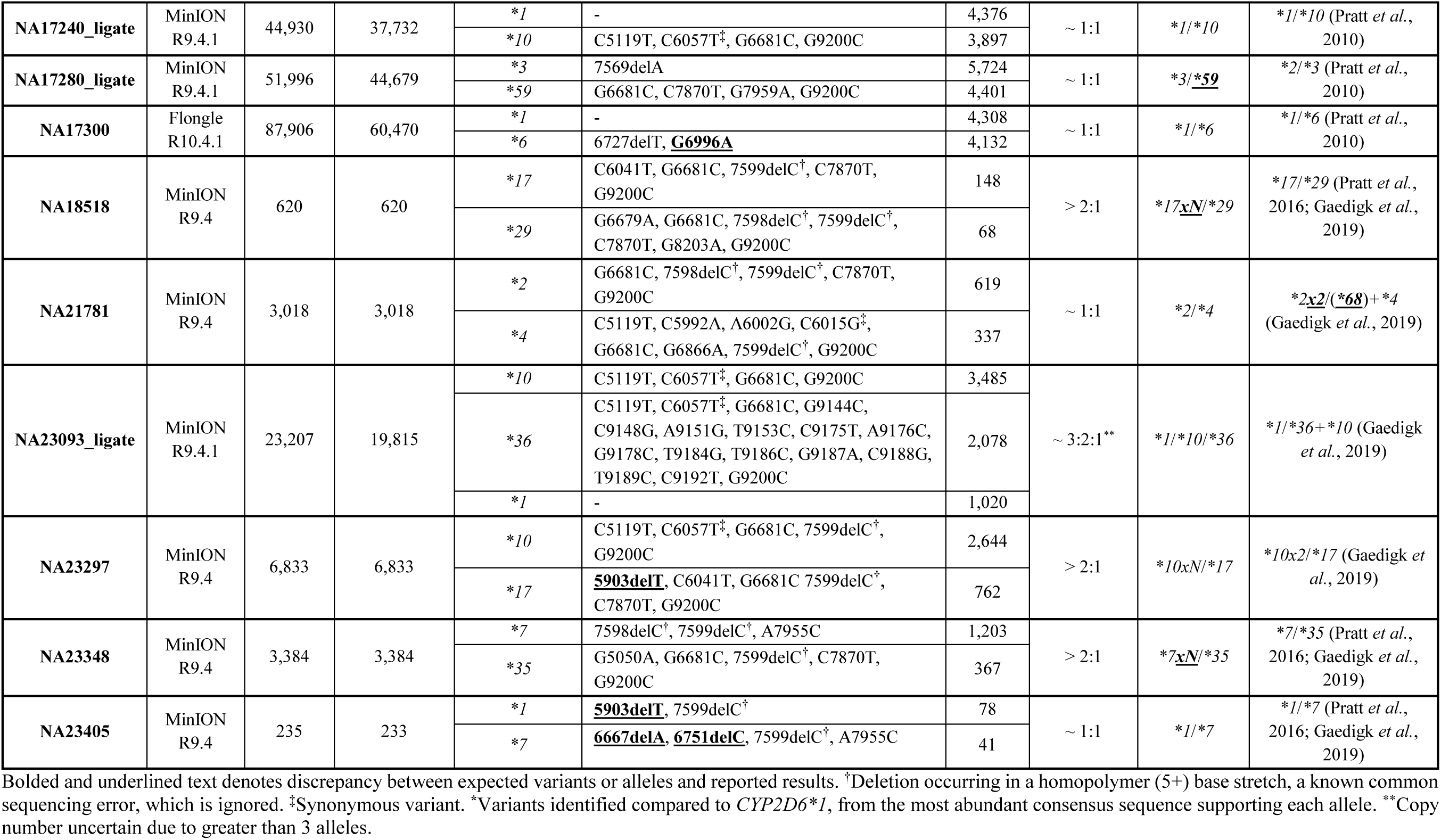
Summary of *CYP2D6* allele reconstruction and diplotyping.

### Inferring copy number from reconstructed alleles

Copy number of genes can be an important factor for determining functional effects. In our results, the relative copy number of the reconstructed alleles could be inferred based on the number of reads that contributed to each allele. We pooled the read counts for all sequences of the same allele. The presence of multiple distinct consensus sequences for the same allele could be used as evidence of duplication events; however, the sequences should be manually inspected to determine if the differences are believed to be from true underlying sequence differences or from regions of poor sequence quality such as those near the ends of the amplicon. Importantly, the lack of multiple distinct sequences does not preclude the presence of a duplication, as each duplicated copy may indeed have an identical underlying sequence. For these reasons, we propose the read depths as a more reliable estimate of relative copy number. To interpret the allele read depth ratios, we tested for ratios > 2:1. In our results, the exact number of duplication events could not be reliably determined, likely due to biases in different allele sequences and sequencing errors, especially in samples with lower read counts. Therefore, the alleles with a greater than 2:1 ratio were assigned an *xN* duplication.

### Known diplotypes are recapitulated from Coriell samples

Across the 22 distinct samples analyzed, our algorithm successfully reconstructed *CYP2D6* diplotypes that matched the reported GeT-RM genotype in 15 cases (68.2 %; **Table 1**) (Pratt *et al*., 2010, 2016; Gaedigk *et al*., 2019). Three cases (13.6 %) did not match GeT-RM, but were consistent with other reports from the literature (described below), suggesting they are also correct. Two cases (9.1 %) only differed by GeT-RM due to copy number differences, and the final two cases (9.1 %) missed the hybrid *\*68* allele which was not detectable due to limitations with the amplicon on those datasets. Overall, this results in an effective 90.9 % accuracy rate when taking into account the current literature and limitations of the amplicons tested. This high level of concordance included 13 samples (59.1 %) containing either synonymous substitutions (C6015G, C6057T) or known homopolymer-induced false-positive deletions (7598delC, 7599delC; **Figure 4A**), which are disregarded during interpretation. This also included seven samples (31.8 %) where allele copy number carried inherent uncertainty in our calls or the GeT-RM reference (NA02016, NA02016_ligate, NA07439, NA07439_ligate, NA18518, NA23297, NA23348). These cases with copy number variation included the two Coriell samples from Wongsurawat et al., where the consensus sequence we generated from their duplication-detecting amplicon supports the presence of a duplicated allele (**Supplementary Figure S3**). In one sample (NA17300) our reconstruction identified an additional uncharacterized variant (G6996A) in a *\*6* allele **(Figure 4B**). This variant occurs downstream of the premature stop codon caused by the 6727delT frameshift mutation, and thus would not have further functional effect. Four uncharacterized deletion variants were detected; 6033delG (NA12878), 5903delT (NA23297, NA23405), 6667delA (NA23405), and 6751delC (NA23405). These all preferentially occurred in consensus sequences with lower numbers of reads, suggesting they may be artefacts due to insufficient coverage rather than true deletions. Inspecting the base support around these regions supports this notion (**Figure 4C-F**), with low coverage in one or both strands.

**Figure 4.**
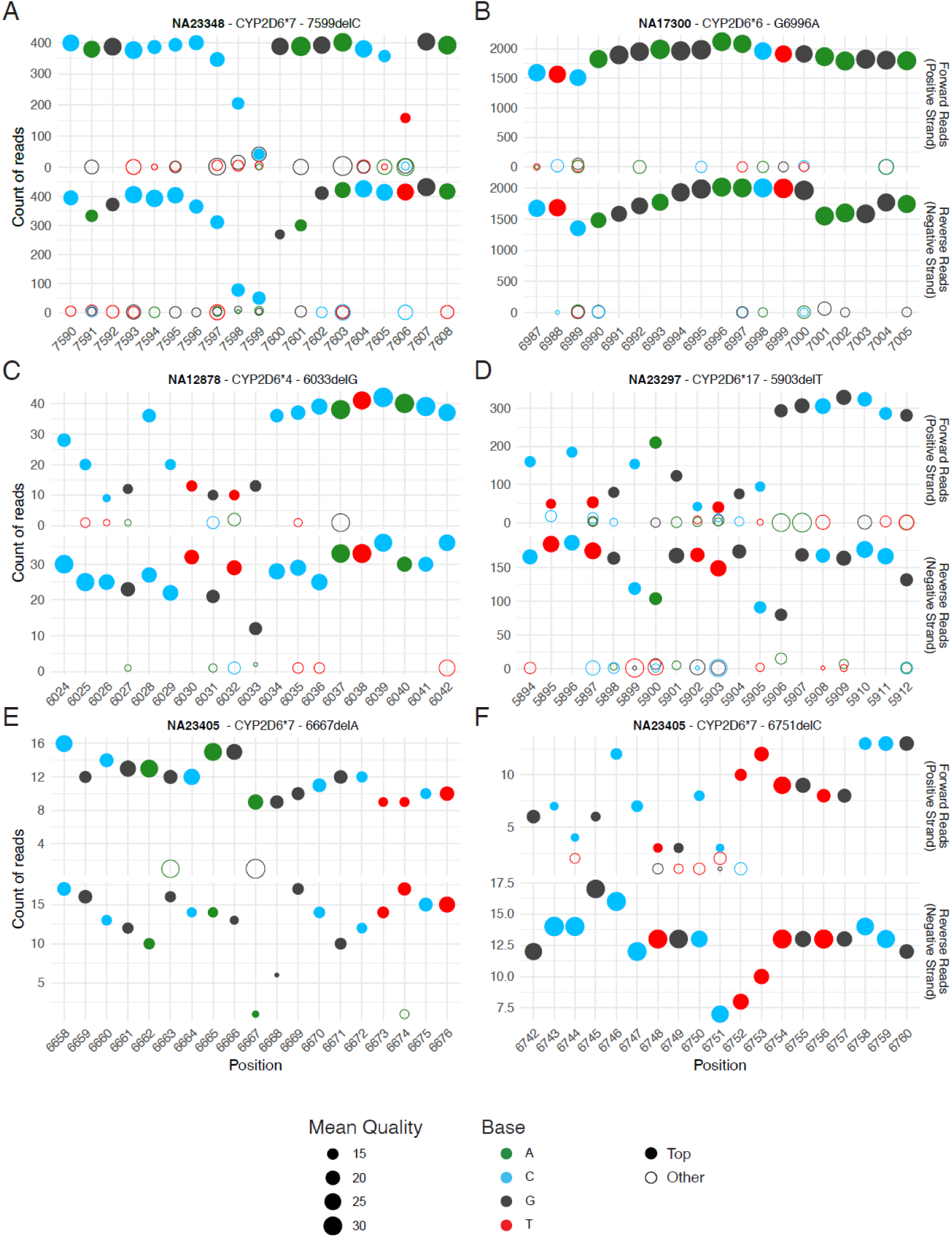
Example signal-to-noise plots for different identified variants. In each panel, the top-observed base at each position is displayed as a solid point, and all other detected bases are hollow points. The size of the point denotes the average quality of the base from all reads used to generate the consensus. The top graph shows results from forward reads, and the bottom graph from reverse reads. (A) An example of the apparent decrease in coverage in homopolymer regions, shown here with a deletion 7599delC being called in NA23348. (B) The novel G6996A variant from NA17300 CYP2D6*6 allele is shown, with no evidence of this being an erroneous call due to poor signal-to-noise or poor quality. This variant occurs downstream of the premature stop codon due to the 6727delT mutation in CYP2D6*6, and thus will not have any effect on the resulting protein. (C-F) Four uncharacterized, non-homopolymer deletions from NA12878, NA23297, and NA23405, all due to low coverage from one or both strands, and are likely technical artefacts rather than true deletions in the sequence.

Of the five samples classified as inconsistent with GeT-RM diplotypes, two (NA17058_ligate, NA17280_ligate) yielded reconstructions strongly suggestive of an error in the reported GeT-RM calls, which were originally based on targeted genotyping panels rather than full-length sequencing, and are known to be imperfect (van der Maas *et al*., 2025) (**Figure 5**). These inconsistent calls are also supported by recent reports (Charnaud *et al*., 2022; Wongsurawat *et al*., 2025). One case (NA17203_ligate) yielded a consensus from *CYP2D7*-derived reads. Recently, NA17203 has been reported as harboring a *CYP2D6*-*CYP2D7* hybrid allele (*\*68*) (Wongsurawat *et al*., 2025; Charnaud *et al*., 2022), though this has not yet been reported by GeT-RM. Manually assessing the alignment of the NA17203_ligate *CYP2D7*-derived consensus sequence to the *CYP2D6* and *CYP2D7* reference sequences supports that this sequence may be representative of the *\*68 CYP2D6*-*CYP2D7* hybrid allele, with a higher number of variants detected from exon 2 – 9 of *CYP2D6* compared to *CYP2D7* (**Supplementary Figure S4**). Indeed, comparing our consensus sequence to the *\*68* sequence obtained by Rubben et al. (Rubben *et al*., 2022) from NA12878, we see only a single indel difference, occurring between exons 3 and 4 (**Supplementary Figure S5**), and thus our consensus sequence supports the same functional hybrid *\*68* allele. The remaining two cases (NA12878 and NA21781) involved the *\*68 CYP2D6-CYP2D7* hybrid allele, which was not detected by the Liau et al. primer design. While no hybrid allele reads were captured in this dataset, the underlying constituent haplotypes were reconstructed correctly, with only the *\*68* classification absent. Taken together, these results demonstrate that the algorithm is robust and largely concordant with reference diplotypes, with nearly all discrepancies attributable to synonymous changes, sequencing artefacts, or structure variants beyond the scope of the amplicon design.

**Figure 5.**
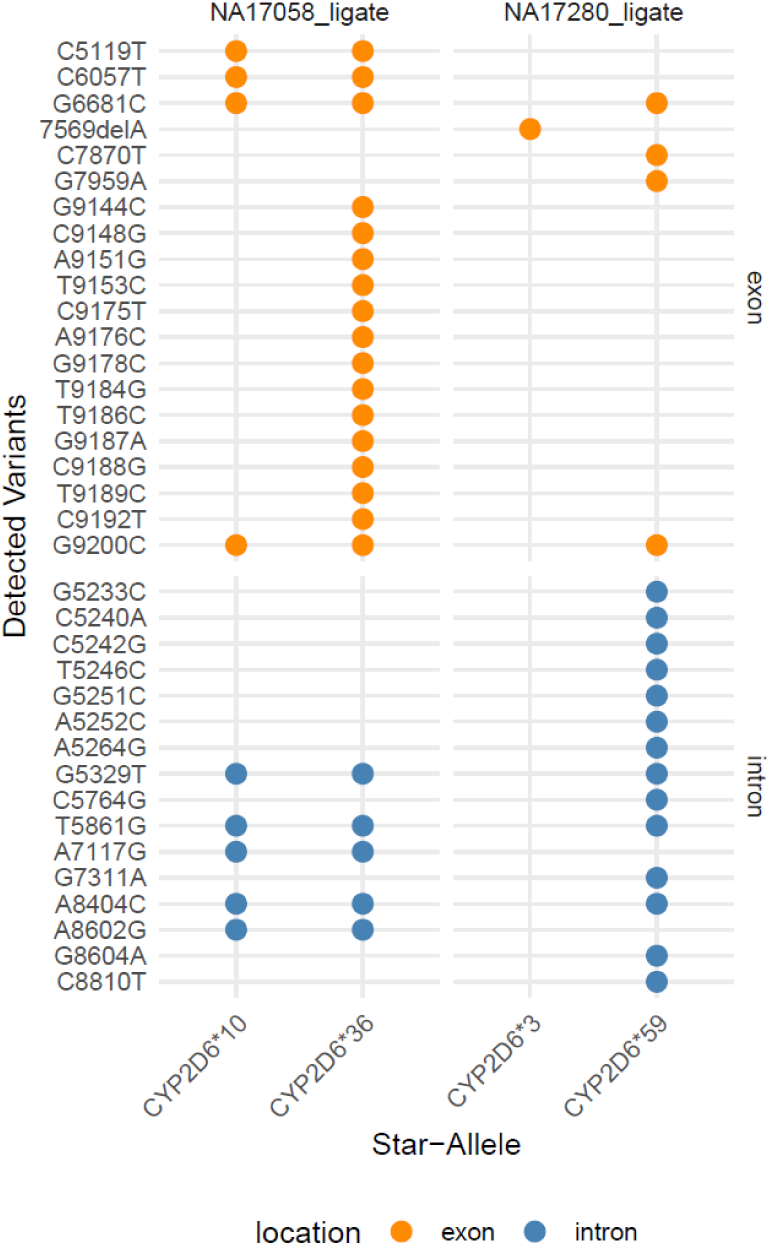
Landscape of variants identified in reconstructed alleles for NA17058_ligate and NA17280_ligate. In each sample, two alleles were reconstructed. Exonic variants are displayed as orange bubbles, intronic variants as blue bubbles. In both samples, the supported alleles differ by a large number of variants, and do not have significant overlap with each other.

### Comparison to existing *CYP2D6* genotyping tools

To put our method in context with existing *CYP2D6* genotyping tools, we ran all of the samples through Aldy4 (Hari *et al*., 2023), SpecImmune (S. Wang *et al*., 2025), and CoLoRGen (Rubben *et al*., 2022), three “variant-first” approaches. SpecImmune and CoLoRGen are compatible with ONT amplicon data. Aldy4 (Hari *et al*., 2023) does not officially support ONT amplicon data, however can be run without phasing or copy number detection to allow genotyping. We did not use any PacBio-specific approaches, as in our experience, ONT and PacBio data and algorithms are generally not cross-compatible due to different sequencing error profiles. For completeness, we also compared our results to the reported genotypes from Liau et al. and Wongsurawat et al. source dataset publications (**Table 2**).

**Table 2.**
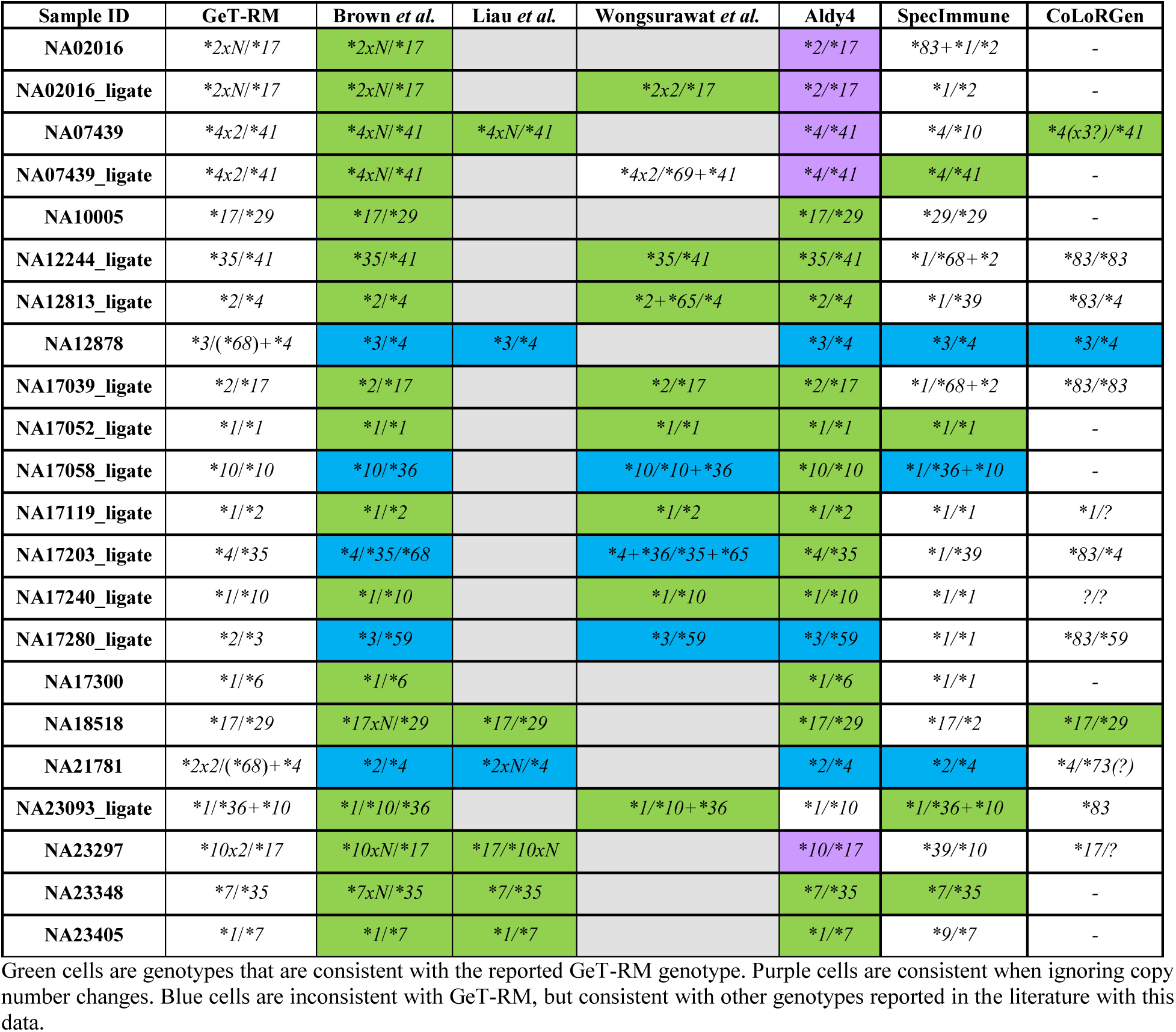
Results from existing *CYP2D6* genotyping tools.

Ignoring copy number inference, Aldy4 generated *CYP2D6* genotypes that matched the GeT-RM genotype in 18/22 (81.8 %) cases (**Table 2**), though we note that two of these are likely incorrect based on more current genotypes available in the literature. Of the remaining four cases, two involved missing a *\*68* allele, and one matched the alternate genotype we and the source publication obtained, further suggesting an error in the GeT-RM genotype. This left one sample with an incorrect genotype call by Aldy4. We obtained *CYP2D6* genotypes for all 22 samples using SpecImmune, though only 5 (22.7 %) matched the reported GeT-RM genotype. Seventeen (77.3 %) had at least one of the two alleles correct. For CoLoRGen, the analysis only completed for 13 samples. Three (13.6 %) matched the GeT-RM genotype, while 8 (36.4 %) had at least one matching allele. Overall, our pipeline demonstrated a greater effective accuracy (90.9 % when considering limitations of the amplicons and current literature; green and blue shading) compared to SpecImmune (31.8 % by the same metric), CoLoRGen (13.6 %), and Aldy4 (86.4 %), while also being able to infer some copy number variations that Aldy4 was unable to do with this data.

### Assessment of another complex region; *HLA*

We extended our analysis to explore three more complex loci; those of the *HLA* class I genes. We identified existing ONT sequence data from 11 subjects who had pooled *HLA-A*, *HLA-B*, and *HLA-C* amplicon data (Anukul *et al*., 2023). Conveniently, this data containing three distinct amplicons also allows us to validate that our pipeline can handle multiple independent loci simultaneously. These datasets had 43,865 ± 6,587 (mean ± standard deviation) reads each. We noted that the coverage of the three loci were unequal; *HLA-A* being covered by 61 ± 12.6 % of the reads, *HLA-B* by 27 ± 11.6 % of the reads, and *HLA-C* by 10 ± 2.8 % of the reads. We ran these 11 samples through our pipeline to reconstruct the allele sequences. To assign the *HLA* allele, we performed a BLAST alignment of each consensus to a database of all *HLA* class I allele sequences obtained from IMGT (Barker *et al*., 2026). For *HLA-A*, we reported the correct two alleles for all samples (**Table 3**). *HLA-B* genotypes had one incorrect allele call in two cases. A single case did not product any *HLA-C* consensus sequence or genotype, likely due to the lower coverage of this locus. Taken together, we show that our allele reconstruction performs well for a second complex locus with sufficient read coverage, with an accuracy of 100 % (*HLA-A*), 90.9 % (*HLA-B*), and 90.9 % (*HLA-C*).

**Table 3.** Summary of HLA genotyping results using data from Anukul et al.

| SRA Run ID | Source Sample ID | HLA Genotype | Our Result |
| --- | --- | --- | --- |
| ERR11404059 | NA02016 | A*30:02:01/A*02:01:01<br>B*53:01:01/B*58:01:01<br>C*04:01:01/C*07:18 | A*30:02:01/A*02:01:01<br>B*53:01:01/B*58:01:01<br>C*04:01:01/C*07:18:01 |
| ERR11404060 | NA07439 | A*29:02:01/A*68:02:01<br>B*42:02:01/B*53:01:01<br>C*04:01:01/C*17:01:01 | A*29:02:01/A*68:02:01<br>B*42:02:01/B*53:01:01<br>C*04:01:01/C*17:01:01 |
| ERR11404061 | NA12244 | A*02:01:01/A*23:01:01<br>B*15:01:01/B*41:01:01<br>C*03:03:01/C*17:01:01 | A*02:01:01/A*23:01:01/<br>B*15:01:01/B*41:01:01<br>C*03:03:01/C*17:01:01 |
| ERR11404062 | NA17039 | A*30:01:01/A*31:01:02<br>B*35:08:01/B*42:02:01<br>C*04:01:01/C*17:01:01 | A*30:01:01/A*31:01:02<br>B*35:08:01/B*42:02:01<br>C*04:01:01/C*17:01:01 |
| ERR11404063 | NA17052 | A*02:01:01/A*33:03:01<br>B*39:04/B*44:03:01<br>C*07:02:01/C*14:03 | A*02:01:01/A*33:03:01/<br>B*44:03:01/-<br>No consensus. |
| ERR11404064 | NA17058 | A*24:02:01/A*26:01:01<br>B*13:01:01/B*40:01:02<br>C*03:04:01/C*08:03:01 | A*24:02:01/A*26:01:01<br>B*13:01:01/B*40:01:02<br>C*03:04:01/C*08:03:01 |
| ERR11404065 | NA17119 | A*03:01:01/A*74:01:01<br>B*35:01:01/B*42:02:01<br>C*07:18/C*17:01:01 | A*03:01:01/A*74:01:01<br>B*35:01:01/B*42:02:01<br>C*07:18:01/C*17:01:01 |
| ERR11404066 | NA17203 | A*24:02:01/A*26:01:01<br>B*14:01:01/B*35:01:01<br>C*04:01:01/C*08:02:01 | A*24:02:01/A*26:01:01<br>B*14:01:01/B*35:01:01<br>C*04:01:01/C*08:02:01 |
| ERR11404067 | NA17240 | A*03:01:01/A*30:02:01<br>B*07:02:01/B*57:01:01<br>C*06:02:01/C*07:02:01 | A*03:01:01/A*30:02:01<br>B*07:02:01/B*57:01:01<br>C*06:02:01/C*07:02:01 |
| ERR11404068 | NA17280 | A*01:01:01/A*11:01:01<br>B*08:01:01/B*18:01:01<br>C*07:01:01/C*07:01:01 | A*01:01:01/A*11:01:01<br>B*08:01:01/B*??<br>C*07:01:01 |
| ERR11404069 | NA23093 | A*11:01:01/A*11:01:01<br>B*15:02:01/B*15:02:01<br>C*08:01:01/C*08:01:01 | A*11:01:01<br>B*15:02:01<br>C*08:01:01 |

The HLA Genotype column provides the known genotypes for these samples, with orange text for those that were missed in Anukul *et al.* (Anukul *et al.*, 2023) (by either their in-house pipeline or Type Stream). Our Result shows the result of our allele reconstruction approach, after using BLAST to identify the best-matching HLA allele. Green text denotes a correct genotype call, while orange text denotes an incorrect call. Our accuracy (percent of 22 alleles correctly identified): HLA-A – 100% (22/22 correct), HLA-B – 90.9% (20/22 correct), HLA-C – 90.9% (20/22 correct).

### Allele reconstruction from other loci

To further test the generalizability of our algorithm to reconstruct allele sequences from ONT amplicon sequence data of any loci, we downloaded representative datasets from SRA. These included two sequence datasets from the *HLA-B* locus (Ton *et al*., 2018), one from *HTT* (Zobel *et al*., 2025), three from *TRAC* (Nitulescu *et al*., 2025), one from *EMX1* (B. Wang *et al*., 2025), one from *CCR5* (Hodge *et al*., 2025), and three from *ACP4* (Liu *et al*., 2026). These datasets had variable sequencing depth (median 22,656.5 reads, range 3,503 – 439,236) and amplicon size (median 1.45 kb, range 289 bp – 9.65 kb). We processed all 11 of these datasets through our pipeline without modification, and compared the resulting consensus sequences with the expected result based on each source publication (**Table 4**). In all but one case, our resulting consensus sequence was consistent with the reported sequence. For data from the *HTT* locus, our resulting consensuses sequence underreported the number of CAG repeats by 7.7 %. Overall, these results demonstrate the utility of our approach for other, unrelated loci.

**Table 4.**
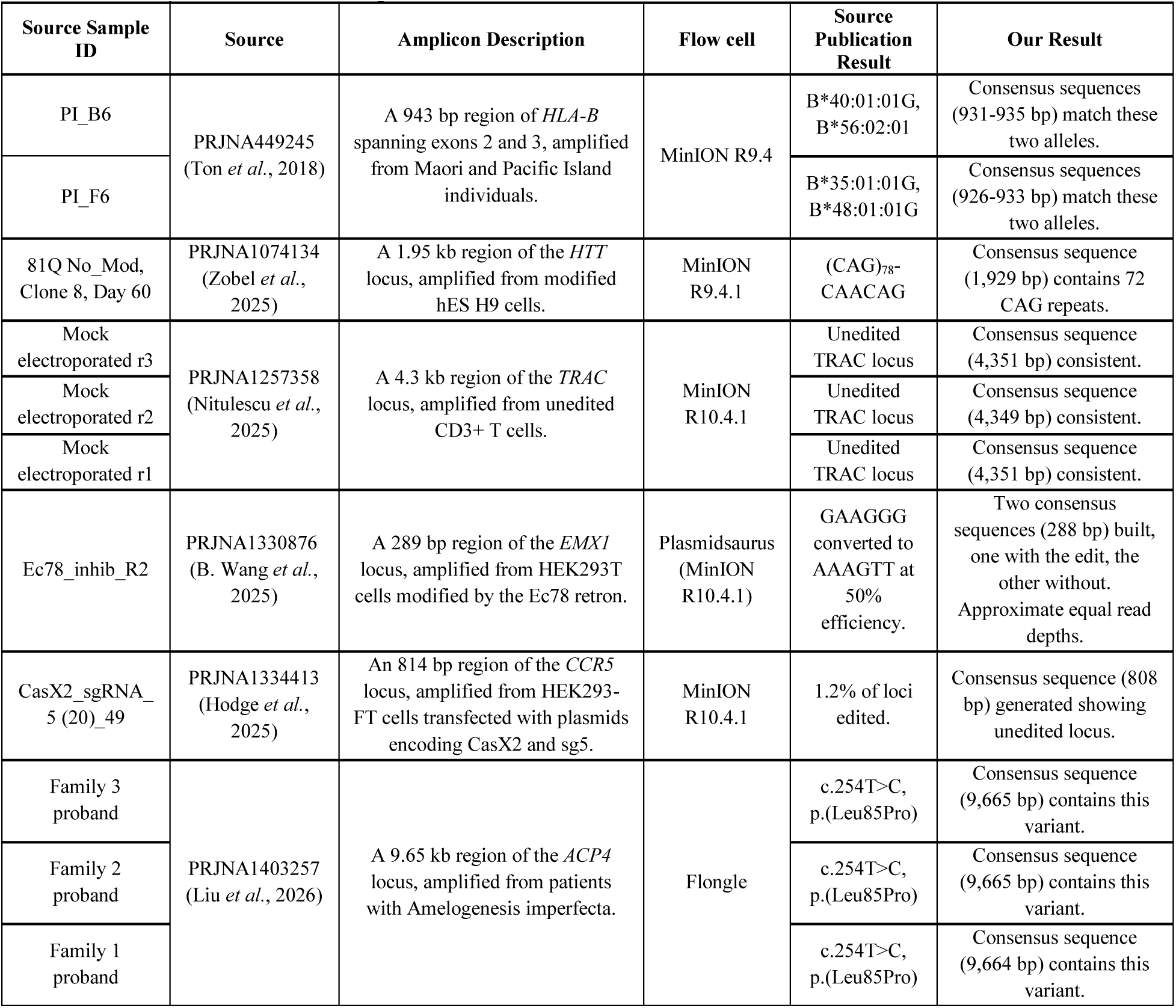
Summary of miscellaneous amplicon reconstruction.

| Source Sample ID | Source | Amplicon Description | Flow cell | Source Publication Result | Our Result |
| --- | --- | --- | --- | --- | --- |
| PI_B6 | PRJNA449245<br>(Ton *et al.*, 2018) | A 943 bp region of *HLA-B* spanning exons 2 and 3, amplified from Maori and Pacific Island individuals. | MinION R9.4 | B*40:01:01G,<br>B*56:02:01 | Consensus sequences (931-935 bp) match these two alleles. |
| PI_F6 |  |  |  | B*35:01:01G,<br>B*48:01:01G | Consensus sequences (926-933 bp) match these two alleles. |
| 81Q No_Mod,<br>Clone 8, Day 60 | PRJNA1074134<br>(Zobel *et al.*, 2025) | A 1.95 kb region of the *HTT* locus, amplified from modified hES H9 cells. | MinION R9.4.1 | (CAG)<sub>78</sub>-<br>CAACAG | Consensus sequence (1,929 bp) contains 72 CAG repeats. |
| Mock electroporated r3 | PRJNA1257358<br>(Nitulescu *et al.*, 2025) | A 4.3 kb region of the *TRAC* locus, amplified from unedited CD3+ T cells. | MinION R10.4.1 | Unedited TRAC locus | Consensus sequence (4,351 bp) consistent. |
| Mock electroporated r2 |  |  |  | Unedited TRAC locus | Consensus sequence (4,349 bp) consistent. |
| Mock electroporated r1 |  |  |  | Unedited TRAC locus | Consensus sequence (4,351 bp) consistent. |
| Ec78_inhib_R2 | PRJNA1330876<br>(B. Wang *et al.*, 2025) | A 289 bp region of the *EMX1* locus, amplified from HEK293T cells modified by the Ec78 retron. | Plasmidsaurus (MinION R10.4.1) | GAAGGG converted to AAAGTT at 50% efficiency. | Two consensus sequences (288 bp) built, one with the edit, the other without. Approximate equal read depths. |
| CasX2_sgRNA_5 (20)_49 | PRJNA1334413<br>(Hodge *et al.*, 2025) | An 814 bp region of the *CCR5* locus, amplified from HEK293-FT cells transfected with plasmids encoding CasX2 and sg5. | MinION R10.4.1 | 1.2% of loci edited. | Consensus sequence (808 bp) generated showing unedited locus. |
| Family 3 proband | PRJNA1403257<br>(Liu *et al.*, 2026) | A 9.65 kb region of the *ACP4* locus, amplified from patients with Amelogenesis imperfecta. | Flongle | c.254T>C,<br>p.(Leu85Pro) | Consensus sequence (9,665 bp) contains this variant. |
| Family 2 proband |  |  |  | c.254T>C,<br>p.(Leu85Pro) | Consensus sequence (9,665 bp) contains this variant. |
| Family 1 proband |  |  |  | c.254T>C,<br>p.(Leu85Pro) | Consensus sequence (9,664 bp) contains this variant. |

To further validate the generalizability of our approach for allele reconstruction, we used our existing sequence data from three plasmids (Brown *et al*., 2023); BCRxV.TF.1 (8.7 kb), BCRxV.VSVG.1 (6.4 kb), and BCRxV.GagPolRev.1 (10.8 kb). Since these plasmids do not occur in the human genome, we built a synthetic genome reference comprised of a single chromosome containing each of the plasmid reference sequences. Upon running these three datasets through our pipeline, we obtained a single consensus sequence for each. All three plasmids resulted in a 99.9 % match to their expected reference. Errors were entirely due to disagreements between strands, resulting in either an N base call or no base being called due to insufficient coverage. We note that this is not unexpected, as our previous work using this data required an alternative pseudo-paired base-calling strategy to resolve strand-specific errors and obtain perfect consensus sequences. Taken together, we have demonstrated the broad applicability of our algorithm to reconstruct the underlying amplicon sequences from amplicon- or ligation-based ONT sequence datasets.

## Discussion

We have presented our solution to reconstruct allele sequences from ONT-based amplicon sequencing, demonstrating its effectiveness on 22 distinct *CYP2D6* sequence datasets encompassing different amplicons and flow cells, 11 pooled *HLA* class I datasets, 11 existing datasets from other genomic loci, and three datasets from plasmid sequencing. Our approach is unbiased, only requiring that a subset of reads exist which fully cover the amplicon of interest, but not requiring *a priori* knowledge of what the specific amplicon is. The appeal of this approach in diverse genes such as *CYP2D6* is that potential duplications and structural variants may be reconstructed without pre-existing knowledge of the sequence. For our application of this approach to the *CYP2D6* locus, we provide post-hoc analysis scripts to automate the variant detection and star-allele assignment, facilitating rapid interpretation of results. Our approach makes no assumptions regarding the number of gene amplicons (or alleles thereof) present in the data, only requiring sufficient coverage of each amplicon in order to be reconstructed and reported. Since the primary output of our approach is the reconstructed allele sequence, novel variants and alleles are readily detectable. In addition to the reconstructed allele sequence and assigned star-allele, our pipeline also outputs a file with per-position base frequencies and average base qualities, allowing interrogation of specific positions to assess the confidence of the resulting base. This gives the user more information, allowing manual interpretation of putative novel variants (**Figure 4**), much like is possible with traditional Sanger sequencing. Within the context of tamoxifen metabolism by CYP2D6, a recent study demonstrated that a neural-network model given full-length gene sequences was better able to predict metabolism rates compared to rates predicted from the assigned star-alleles alone (Van der Lee *et al*., 2021). This demonstrates the increased power of allele sequence reconstruction, and suggests that such reconstructions may be even more important in the future as more machine learning models are trained across a wider range of genes and pathways.

The concept of building a consensus from clustered reads is not unique, however, our approach differs from existing methods in the following ways. Buermans *et al*. describe the application of the Pacific Biosciences long-amplicon analysis tool (pblaa) to *CYP2D6* (Buermans *et al*., 2017), an approach that is based upon the concept that multiple unique consensus sequences are representative of multiple alleles. This approach is specific to PacBio sequence data, and is closed-source. Athlon (Liu *et al*., 2018) is a tool developed to *HLA* type ONT data, performing an alignment to *HLA* reference alleles to partition reads, and variant calling to build the consensus sequences. This is limited to two alleles per *HLA* gene, requires upfront segmentation of reads into specific loci, and only uses a subset of total reads. Hui *et al*. (Hui *et al*., 2025) describe an approach to generate consensus 16-23S rRNA sequences for microbiome analysis. This approach orients sequence reads based on known primer sequences, clusters reads by similarity, and generates a consensus using Racon or Medaka. While this approach performs well for 16S genes, it is unable to distinguish separate alleles of human genes differing by a small number of variants, such as SNVs, since reads from both alleles would be clustered together, and variants would be “error-corrected” away. Similarly, Sahlin *et al*. (Sahlin *et al*., 2021) describe a cluster-consensus approach for use in the microbiome field, which suffers from the same limitations when applied to highly similar alleles of human genes, lacking the sensitivity to output separate consensus sequences that may legitimately differ by only a single base.

We observed a higher than expected number of short reads in our data. In assessing these reads after mapping to the genome, they appear to be spread across the whole expected *CYP2D6* amplicon region, and are not coming from any single contaminant (**Supplementary Figure S1**). Such a signal is consistent with the concept of DNA amplicon fragmentation during sequence library preparation, resulting in these DNA fragments having adapters ligated and subsequently sequenced. We note that Liau et al. (Liau, Maggo, *et al*., 2019) also observed a large fraction of short reads in their data (> 90 % of their reads were shorter than the expected ∼6 kb amplicon in their first sequencing run, ∼45 % in their second run after changing the beads used for size selection to remove DNA fragments shorter than 3-4 kb), so this may be an expected artefact of sequence library preparation which would require handling in the downstream analysis pipeline. We opted to only consider short reads if they were evidenced to have been derived from a single conserved, short amplicon.

Existing methods for diplotyping *CYP2D6* from amplicon or whole genome sequence data take a “variant-first” approach (Hari *et al*., 2023; Chen *et al*., 2021; Deserranno *et al*., 2023; Liau, Maggo, *et al*., 2019; Ammar *et al*., 2015), using algorithms such as Clair3 (Zheng *et al*., 2022) to call variants, and then these variants are subsequently phased into haplotypes using tools like WhatsHap (Patterson *et al*., 2015). Our method differs from existing methods in that it is not reliant on variant calling directly from sequence reads. Rather, we distil the set of sequence reads into high-confidence, reconstructed allele sequences, which can be directly compared to representative reference allele sequences for haplotype determination. This is especially helpful when analyzing ONT reads, which have higher rates of sequencing errors than NGS approaches. Our allele sequence reconstruction approach is not reliant on a specific number or identity of amplicons within the data, theoretically handling even extreme cases of copy number variation or gene duplication. Our method uses commonly available Linux tools and open-source software, defined in Anaconda environments and should be readily deployable on any Linux system.

We ran three existing *CYP2D6* diplotyping tools on our datasets; Aldy4, SpecImmune, and CoLoRGen. Aldy4, despite not officially supporting ONT data, had similar accuracy as our method. Unfortunately, copy number detection was unable to be performed by Aldy4 for amplicon data, and so results from Aldy4 do not capture the gene duplication events that our method was able to identify. SpecImmune generally had lower accuracy compared to our method, with the reported genotype matching the GeT-RM genotype in only five samples. CoLoRGen similarly had lower accuracy, with only three samples matching the GeT-RM genotype. Further, CoLoRGen failed to generate a genotype for nine samples.

Our approach requires amplicon data, and specifically requires reads to be uniform in their coverage of the amplicon. As such, the ONT Rapid (transposase-based) library construction kits are incompatible, as are “shotgun” based approaches such as ONT adaptive sampling, as they result in read sets with variable start and end positions. Further, with data from a single amplicon, there is no way to unambiguously assess copy number. Based on the number of reads contributing to each reconstructed allele, we can approximate the copy number ratio if a sample has two distinct allele sequences. We note that this is a relative abundance measure rather than an absolute measure of copy number. Despite this, relative abundance is likely sufficient to ascertain clinical effect, as all *CYP2D6* alleles capable of ultrarapid metabolism do so by duplicating at least once (Hicks *et al*., 2017). Thus, determining the relative abundance of normal functioning alleles may be sufficient to designate an ultrarapid metabolizer. Further copy number increases will elevate the “Activity Score” of the diplotype, but will not change the metabolizer status, nor drug prescription recommendations. On the other hand, since there is no genomic reference to accurately estimate relative copy number, if certain diplotypes consist of duplicated homozygous normal function alleles (such as *\*1*/*\*1×2*, *\*2*/*\*2×2*, *\*33*/*\*33×2*, etc.), it is impossible to determine the copy number and thus the metabolizer phenotype. Multiplexing of multiple genes may help, assuming that one locus is copy number invariant. Alternatively, a control region of the genome could be included that is known to be copy number invariant, as is done for data used by Aldy 4 (Hari *et al*., 2023). The data from Wongsurawat et al. (Wongsurawat *et al*., 2025) demonstrated that our analysis of *CYP2D6* is not hindered by the presence of multiple amplicons, supporting this as a reasonable approach in the future.

For the 22 *CYP2D6* datasets examined in this manuscript, none contained any detectable population of reads that contained full-length gene duplication events, likely due to the standard *CYP2D6* primers resulting in too large of an amplicon to be efficiently amplified in the event of a gene duplication. Wongsurawat et al. included specific primers targeting the region between duplicated gene copies, which produced a readily identifiable 3.5 kb amplicon in their two samples with known gene duplications. Two samples from Liau et al. with known *CYP2D6*68* alleles (a gene hybrid with *CYP2D7*) were both called incorrectly, with no reads showing the expected fusion event. Alternative primer sets would be worth exploring in the future to increase the chances of capturing these structural changes and other complex tandem rearrangements, which have not yet been tested using our analysis approach. Gene deletions, such as *CYP2D6*5*, are also detectable by our pipeline as long as a sufficient population of reads exist which span the deleted region. We validated this using simulated reads, demonstrating that reads spanning a large deletion in the genomic reference sequence are effectively reconstructed into the true, deletion containing consensus sequence. Overall, while we have not comprehensively tested the entire landscape of all known *CYP2D6* variants, we show consistently positive performance reconstructing the underlying allele sequences for 14 distinct alleles from data sets for which there are reads supporting these alleles.

Our method is extensible to other genomic loci. We ran our allele reconstruction on 11 samples containing pooled data from *HLA-A*, *-B*, and *-C*, and show above 90 % accuracy in the resulting HLA genotypes. This supports the use of this method for other complex loci in the human genome. More generally, we identified 11 existing ONT sequence datasets from six different genomic loci, demonstrating that our approach faithfully reconstructs the underlying amplicon sequence in each case. Finally, we show that this approach can be applied to any amplicon-like sequence where an expected reference sequence is known, running it on plasmid sequence reads for 3 different plasmids. The reconstructed plasmid sequences showed 99.9 % accuracy.

We have demonstrated that *CYP2D6* allele sequence reconstruction and diplotyping can be achieved from modest long-read sequencing (ONT Flongles), requiring limited specialized equipment, and having low cost. It is appealing that a singular sequencing reaction coupled with this algorithm may be able to ascertain *CYP2D6* diplotypes with accurate phasing of known or novel variants, as well as detect gene duplications and structural variants without the need for multiple experiments. This information can then be used to designate metabolizer status in order to facilitate rational drug prescription as per CPIC guidelines (Hicks *et al*., 2017). Currently, to achieve this clinically, a laborious series of assays are conducted, such as Taqman real-time PCR reactions for common SNPs and indels, copy number verification by qPCR and finally long-range PCR is used for confirmation of duplications or deletions (Pratt *et al*., 2021). These assays must have regulatory approval, being performed in a qualified lab using approved medical devices. Regulatory approval of our approach is possible, particularly by transitioning to the GMP-ready Q-Line of ONT devices. This would necessitate, at the expense of more complexity, barcoding and multiplexing of samples on larger MinION flow cells to increase throughput.

## Materials and Methods

### CYP2D6 gene amplification

Genomic DNA was quantified using the Nanodrop, and diluted to 200 ng of DNA in 33.5 uL of nuclease-free water. Primer sequences were obtained from Ammar et al. (Ammar *et al*., 2015), using CYP2D6-2F (TAGCTCCCTGACGCCATGATTTGTCTT) and CYP2D6-2R (CCTGGTTATCCCAGAAGGCTTTGCAG). We used the standard protocol for Phusion High-Fidelity DNA Polymerase (NEB M0530): 5× Phusion HF Buffer, 10 mM dNTPs, 10 uM forward primer, 10 uM reverse primer, 200 ng template genomic DNA, and 0.5 uL Phusion DNA polymerase in a 50 uL reaction. PCR cycling conditions were: 98°C for 30 sec., followed by 35 cycles of 98°C for 10 sec. and 72°C for 120 sec., followed by a final step of 72°C for 10 min. and hold at 4°C. PCR products were cleaned up using the Qiagen QIAquick PCR Purification Kit following manufacturer’s instructions.

### Sequence library construction

Libraries were prepared using the ONT Ligation Sequencing Kit V14 (SQK-LSK114) following the provided protocol (LSK114_ACDE_9163_v114_revN_29Jun2022). Samples were quantified with Qubit dsDNA HS Assay kit (Invitrogen Q32851). One hundred fmols of amplicon DNA was repaired using NEBNext Ultra II End Prep Enzyme Mix provided in the NEBNext Companion Module for Oxford Nanopore Technologies Ligation Sequencing kit (NEB E7180S). Samples were purified with 30 uL of Ampure XP beads (Beckman, A63880), and eluted with 31 µL of nuclease-free water. Adapter ligation was performed with the NEBNext Quick T4 DNA Ligase provided in the NEBNext Companion Module, ONT Ligation Buffer (LNB) and ONT adapter mix (LA). Samples were purified with 20 uL Ampure XP beads, washed with ONT Long Fragment Buffer, and eluted with 7 µL ONT Elution Buffer (included in SQK-LSK114 kit).

### Sequencing using ONT MinION

The Flongle R10.4.1 (FLO-FLG114) flow cell was placed into the Flongle Adapter provided with the Flongle Starter Pack (FLGIntSP). Prior to sequencing samples, flow cell check was performed following the ‘flow-cellpcheck-PQE_1004-V1’ protocol to ensure a minimum of 50 active pores were present. The flow cell was prepared and run according to ONT instructions: the recommended 5-10 fmol of prepared library was loaded per flow cell, and the run time was set to 24 hours. The Flongle Adapter was USB connected to an M1 Macbook Pro with 16 GB of RAM. On-device base calling was disabled, instead performing base calling in a later step on our higher-powered computational resources.

### Base calling raw sequence data with Guppy

We assembled all steps of our pipeline into a Snakemake v6.3.0 workflow. We performed initial base calling of the raw .fast5 data using the GPU version of Guppy v6.5.7, selecting the “dna_r10.4.1_e8.2_400bps_sup.cfg” base calling configuration file. We ran base calling on a GPU cluster comprised of eight NVIDIA GeForce RTX 3090 GPUs, 256 GB RAM, and two Intel Xeon Silver 4212 12 core CPUs. All .fast5 files generated by MinKNOW for each sample were submitted to the cluster in its own job, requesting 20 GB RAM, 1 GPU, and 4 CPUs. To obtain the most useable data possible, we retain the failed reads, as reads with poor quality at critical positions will be filtered out later in our pipeline. We ran Porechop v0.2.4 (https://github.com/rrwick/Porechop) to remove any remaining adapter sequences and separate chimeric reads.

### Independent control data

We obtained existing base called sequence data of *CYP2D6* from other Coriell samples from Liau et al. (Liau, Maggo, *et al*., 2019) and Wongsurawat et al. (Wongsurawat *et al*., 2025). These datasets were generated in different labs, using different primers, and sequenced on different versions of the ONT flow cells. These datasets are used to help us validate that our process is not dependent on a specific amplicon or data source. Class I *HLA* data from Anukul et al. (Anukul *et al*., 2023) was downloaded to test the performance on a different complex locus, and to test the performance on datasets with pooled amplicons from multiple independent loci. Data from other human genomic loci were obtained for the *HLA-B* locus (Ton *et al*., 2018), the *HTT* locus (Zobel *et al*., 2025), the *TRAC* locus (Nitulescu *et al*., 2025), the *EMX1* locus (B. Wang *et al*., 2025), the *CCR5* locus (Hodge *et al*., 2025), and the *ACP4* locus (Liu *et al*., 2026). Data from plasmid sequencing of three plasmids was obtained from Brown et al. (Brown *et al*., 2023), using the subset of reads the authors made available on Zenodo.

### Inference of genomic amplicon

All of these raw reads are aligned to the human genome (hg38) using minimap2 v2.17-r941 (Li, 2018) with the following parameters: “-x map-ont --secondary=no –t 4”. To infer the reference amplicon sequence in an unbiased manner, we use samtools v1.15 (Danecek *et al*., 2021) to get the depth of read coverage at every position of the genome, and extract out the regions that have depth above 10 % of the max observed depth. This parameter is tunable, potentially needing to be lowered for datasets comprised of amplicons from multiple distinct loci with uneven coverage. A helper script is made available on the GitHub code repository to visualize the distribution of depths to help select an appropriate threshold if needed. Any adjacent regions are merged using bedtools v2.26.0 (Quinlan and Hall, 2010) with the following parameters: “merge -d 100”. To aid in the detection of amplicons supporting large deletions, we check all pairs of high-coverage regions for reads which align to both regions. If 50 % of the reads aligning to each region are shared, we create “chimeric regions” made by concatenating these regions, which are used in addition to each region alone in subsequent steps.

### Candidate read extraction

We extract all the reads that had a primary or supplemental alignment to the identified genomic region(s). We observed a high number of short reads in our in-house-generated data. To further group reads into those completely covering the same amplicon sequence, we perform length clustering on the candidate reads. Clustering is done using the DBSCAN algorithm (Schubert *et al*., 2017) from the scikit-learn library (v1.8.0), run on the read lengths, with the allowed neighbour length variability (EPS) set to 10 and the minimum core point neighbours set to 0.1% of the total number of candidate reads, or 5, whichever is greater. This ensures that data sets with fewer reads still have a reasonable value used. We performed a brief parameter space exploration on three samples, testing all pairwise combinations of EPS values 10, 20, 30, 40, and 50, and minimum neighbor values 5, 10, 15, 25, and 50. The maximum number of reads in the productive length cluster in each case was obtained with EPS 50 and minimum neighbours 5, however, these large clusters had highly variable read lengths. Decreasing EPS to 10 and increasing minimum neighbours to 50 dropped the number of reads by an average of 41.3 %, but resulted in clusters of reads with highly similar lengths; optimal for consensus generation. However, we noted in small data sets, a minimum neighbour value of 50 was too stringent, thus we implemented the value to scale with the input number of reads. Clusters are selected for consideration for downstream allele reconstruction if the number of reads in the cluster is at least 25 % of the size of the largest cluster, selected empirically. Additionally, to ensure we do not miss the cluster with reads the full-length of our inferred amplicon, we select the cluster with the length closest to the inferred amplicon region if it is not already included.

### Allele sequence reconstruction

Our allele reconstruction method is built as a Python (v3.11) script, and takes the alignments of each group of candidate reads to their inferred amplicon reference sequence obtained using minimap2 v2.17-r941. These alignments are parsed, along with the raw fastq files to extract per-base quality scores and unaligned sequence flanking each alignment. The unaligned flanking sequences are treated as insertions, and are only used if at least 25% of the reads support sequence at that position. To facilitate efficient assessment of the pile-up of base support at every reference position and possible insertion sites, we use byte matrices to store a representation of the multiple sequence alignment of all reads. To generate this, we step through every alignment to track the maximum observed insertion at every position. We then create empty byte matrices of reads × positions, where positions are offset relative to the reference sequence to allow for all observed insertions. We then step through all alignments again to load the observed bases and base quality scores into the byte matrices. The element i,j of the matrix are integer representations of the base found in the ith read at position j. Separate matrices are maintained for positive and negative strand alignments, facilitating strand-specific assessment of coverage. The matrices are evaluated, position by position, to assess the number of reads supporting each base at each position. If a position has evidence of heterozygosity (here, defined as the top two most-observed bases having less than a 2:1 ratio in abundance, threshold determined empirically), the reads are partitioned into separate groups based on support of the heterozygous bases (**Figure 3C-D**), and consensus sequence construction tries again on each of the new read groups independently. This repeats, recursively, until pure (no evidence of heterozygosity) consensus sequence(s) are derived, or the read groups are too small to continue. Due to the phenomenon of strand-specific errors in ONT data (Krishnakumar *et al*., 2018; Branton *et al*., 2008), this test of purity is also performed on the forward and reverse reads independently. Based on a 5kb reference sequence, our algorithm requires 1 CPU core and approximately 3 GB of RAM per 10,000 reads.

, The allele reconstruction script has a number of parameters that can be adjusted to fine-tune the behavior of the allele reconstruction. Based on all of the control data, we used the following settings, determined empirically, and explained subsequently: --min_depth_factor 0.25, --absolute_min_depth_thresh 10, --min_base_qual_global 10, --min_base_qual_percentile 33, --sig_to_noise_thresh 2. The “--min_depth_factor” controls the minimum number of reads needed to support a base at any position, and represents the fraction of the maximum observed depth (here, 25 %). The “--absolute_min_depth_thresh” controls the minimum number of reads supporting a reconstructed allele sequence for it to be output, here set to 10 reads. The “--min_base_qual_global” is the minimum base quality score needed to be used for allele sequence reconstruction. If this threshold is not met, then the “--min_base_qual_percentile” is used, being a percentile threshold, per position, that is the lower base quality limit. This allows for data to be obtained from positions with lower quality overall. Finally, the “--signal_to_noise_thresh” is the value of the minimum ratio of the most frequent base to the second most frequent base at any position needed to be considered homozygous. This needs to be met when considering all reads, and is multiplied by a factor of 1.5 when also testing reads from each strand separately. In the case of a position appearing heterozygous when taking all reads together, but not when assessing each strand individually, the top observed base from the pooled strands is tested against a softened signal to noise threshold (divided by a factor of 1.5) to call a consensus base, or an N base if a consensus base cannot be determined. Based on quality thresholds, a read may be “ambiguous” for a given position (supporting a base, but due to low quality the base is not used). Ambiguous reads do not contribute to depth calculations at that position. If the position is heterozygous, ambiguous reads for that position are discarded. The average quality score for each consensus base across all reads used for the consensus is recorded and output in the detailed results file, along with base frequencies, allowing manual inspection of the reconstructed allele sequences.

### Automatic genotyping of consensus sequence

Each resulting reconstructed allele sequence is aligned to the *CYP2D6* reference sequence using Biopython v1.79 pairwise2.align.localxs, and the alignment is parsed to enumerate all variants. All known *CYP2D6* variants are catalogued from the *CYP2D6* Allele Definition Table from ClinPGx (Whirl-Carrillo *et al*., 2021, 2012), downloaded from https://www.pharmgkb.org/page/cyp2d6RefMaterials on January 21, 2025 (now available at https://www.clinpgx.org/page/cyp2d6RefMaterials). This file was manually curated to ensure consistency in how each variant type was reported, allowing for straightforward parsing. For variants that can have multiple possible bases allowed for a given allele, all possible allowed values are stored. The set of identified variants for each reconstructed allele sequence is compared to those from each known allele, and a score is calculated for each known allele based on the number of disagreements in identified variants within the exon and splice regions of the gene. The allele with the lowest score is returned as the best candidate allele assignment for the reconstructed allele sequence, along with any missing or extra variants present in the consensus sequence compared to the star allele reference should they exist. In this way, manual review of the candidate alleles and identified variants allows a user to confirm the allele assignment, or characterize a novel allele. A helper script is made available on the GitHub code repository to report out the most likely genotype for each sample based on all identified consensus sequences and abundances. This combines counts for all consensus sequences supporting the same *CYP2D6* allele, and outputs any allele that represents at least 5 % of the reads of that length-cluster. Alleles with a greater than 2:1 ratio to each other are reported as possibly supporting a duplication or copy number variation event.

### Benchmarking to Aldy4, SpecImmune, and CoLoRGen

We directly compared our results for the 22 *CYP2D6* datasets to results we obtained from running Aldy4 (Hari *et al*., 2023), SpecImmune (S. Wang *et al*., 2025), and CoLoRGen (Rubben *et al*., 2022), three “variant-first” approaches. Aldy4 was installed as described on the GitHub repository (https://github.com/0xTCG/aldy) using pip to install the required packages into a conda environment. To allow Aldy4 to run on ONT amplicon data, copy number detection and variant phasing needed to be disabled. Aldy genotype was run with “--gene cyp2d6 --profile illumina -o [samplename]-cyp2d6.aldy --cn 1,1 input.bam --param sam_long_reads=true phase=False”. We installed SpecImmune as described on the GitHub repository (https://github.com/deepomicslab/SpecImmune) using conda, after simplifying the provided conda environment snapshot file to be compatible with our system. SpecImmune was run with “-r porechopped-reads.fastq -j 16 -i CYP -n samplename -o output_directory -y nanopore --db CYP_ref_db --hg38 hg38_ref.fa”. We used the [samplename].CYP.merge.type.result.txt result file for each sample, taking the reported “CYP2D6 Diplotype”. CoLoRGen was installed from the GitHub repository (https://github.com/laurentijntilleman/CoLoRGen), creating a conda environment with python 3.8, medaka 1.4.3, pandas 1.1.5, gtfparse 1.2.1, samtools 1.11, minimap2 2.18, tabix 1.11, py-bgzip, bowtie2, and pip to install bio 1.5.9. After manual adjustment of the provided parameter file, removing unnecessary fields, we ran CoLoRGen with the provided parameters. CoLoRGen failed on nine samples for unknown reasons. We used the haplotypes_haplotypes.txt file for the final genotype calls.

### Simulated deletion detection

To test our algorithm’s ability to reconstruct alleles that represent a large deletion relative to the reference genome, we simulated ONT reads using PBSIM3 (v3.0.5; https://github.com/yukiteruono/pbsim3). We provided a reference template sequence derived from our amplicon sequence with the entire 4.3 kb *CYP2D6* gene deleted, resulting in an 836 bp template. To simulate 1,000 reads, we created a template reference fasta file containing this same sequence as 1,000 separate entries. We ran PBSIM3 using qshmm QSHMM-ONT-HQ.model, downloaded from the GitHub repository. Upon running our pipeline on these simulated reads, we obtained a consensus sequence built from 910 reads which exactly matched the provided template sequence.

### Testing performance on HLA class I sequence data

We ran our pipeline with the following modifications to test our ability to generate *HLA* consensus sequences from the data from Anukul et al. First, after inspecting the relative depths of the *HLA-A*, *HLA-B*, and *HLA-C* loci in these samples, we lowered the LOCI_MIN_COVERAGE parameter from 0.10 to 0.04 to ensure that all *HLA* loci were retained. In this data, the coverage from *HLA-C* is as low as ∼4 % of the coverage of *HLA-A* in some samples. Second, to make *HLA* genotype calls on the resulting consensus sequence, we replaced our automated *CYP2D6* genotyping steps with a BLAST alignment (blastn v2.5.0) (Camacho *et al*., 2009) to a database of *HLA* class I allele sequences downloaded from IMGT (Release 3.63; 2026-01) (Barker *et al*., 2026), using the following parameters: “-word_size 11 –reward 2 –penalty -3 –gapopen 5 –gapextend 2 –dust no –soft-masking false –evalue 1e10 –max_target_seqs 50”. The top scoring *HLA* allele(s) were output as the best match for each consensus sequence, with genotypes collapsed at the 6-digit level.

### Testing performance on other genomic loci

When testing our pipeline on sequence datasets from other, non-complex loci, we ran our pipeline without modification, aside from skipping the final *CYP2D6* variant calling and allele assignment steps. Resulting consensus sequences were manually assessed based on the reported results from each source publication, checking if our consensus sequences were consistent with the results already reported.

### Testing performance on plasmid sequence

To test our algorithm’s ability to reconstruct sequence not derived from the human genome, we used our existing plasmid sequence data. To perform this analysis, we created an appropriate reference sequence comprised of a single chromosome containing the reference sequences of our plasmids. All other aspects of the pipeline were left unchanged, except for skipping of the final *CYP2D6* variant calling and allele assignment. Resulting consensus sequences were compared directly to the known plasmid sequence for characterization of any sequence errors. Importantly here, we did not perform the pseudo-paired basecalling as described in our previous work (Brown *et al*., 2023), which was required to obtain perfect sequence verification in these plasmids. We chose to analyze these sequences in a more standard manner here, to better represent typical workflows.

## List of Abbreviations

CPU: Central Processing Unit
CYP: Cytochrome P450
DNA: Deoxyribonucleic Acid
GPU: Graphics Processing Unit
HLA: Human Leukocyte Antigen
NGS: Next Generation Sequencing
ONT: Oxford Nanopore Technologies
PCR: Polymerase Chain Reaction

## Declarations

### Ethics approval and consent to participate

This study was approved by the Fraser Health Research Ethics Board and BC Cancer Agency Research Ethics Board (H21-01422).

### Consent for publication

Not applicable.

### Availability of data and materials

Custom code is available at the following GitHub repository, along with instructions for use and test data: https://github.com/scottdbrown/allele-reconstruction-long-read-amplicon-data. A snapshot of the code at the time of publication is available on Zenodo.org; doi 10.5281/zenodo.19716004. Raw .fastq sequence data for our three sequencing runs is available at the SRA under Bioproject PRJNA1357883 (https://www.ncbi.nlm.nih.gov/bioproject/1357883).

### Authors’ Contributions

RAH, PIM, and SDB designed the study. LD performed library preparation and sequencing optimization. SM, AL, MK, AC, and MM performed library preparation and sequencing. AM, NW, and MH coordinated and oversaw the technical work conducted in the Molecular Cytogenetics Laboratory at Royal Columbian Hospital. SDB performed the sequence analysis and algorithm development. SDB and PIM wrote the manuscript. All authors read and approved the final manuscript.

## Acknowledgments

We would like to thank Dr. Martin Kennedy for sharing additional data from Liau et al. (Liau, Maggo, *et al*., 2019).

## Supplemental

**Supplementary Table S1.** Summary table of all inferred amplicons, length-clusters, and allele reconstruction attempts with read counts at each stage.

**Supplementary File S1.** All successfully constructed consensus sequences for all *CYP2D6* samples.

**Supplementary Figure S1.**
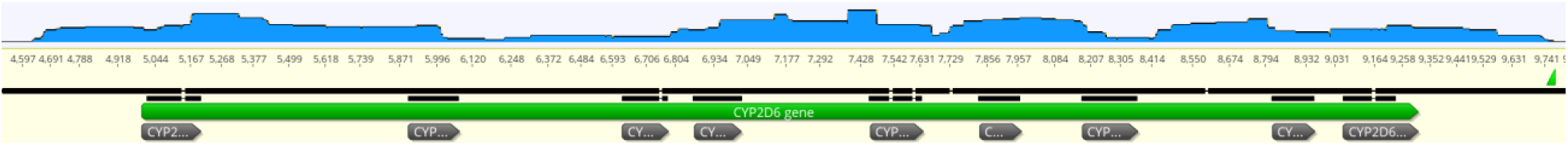
*CYP2D6* read coverage from short reads. Screenshot from Geneious Prime. The length-cluster of short reads from NA17300 (40,717 reads with mean length 445 bases) was aligned to the *CYP2D6* reference sequence. Coverage is shown in blue, spanning the entire expected amplicon. Exons are marked with grey segments under the green *CYP2D6* gene. Coverage is variable, and the region of highest coverage (exon 4-6) yielded a single reconstructed consensus sequence.

**Supplementary Figure S2.**
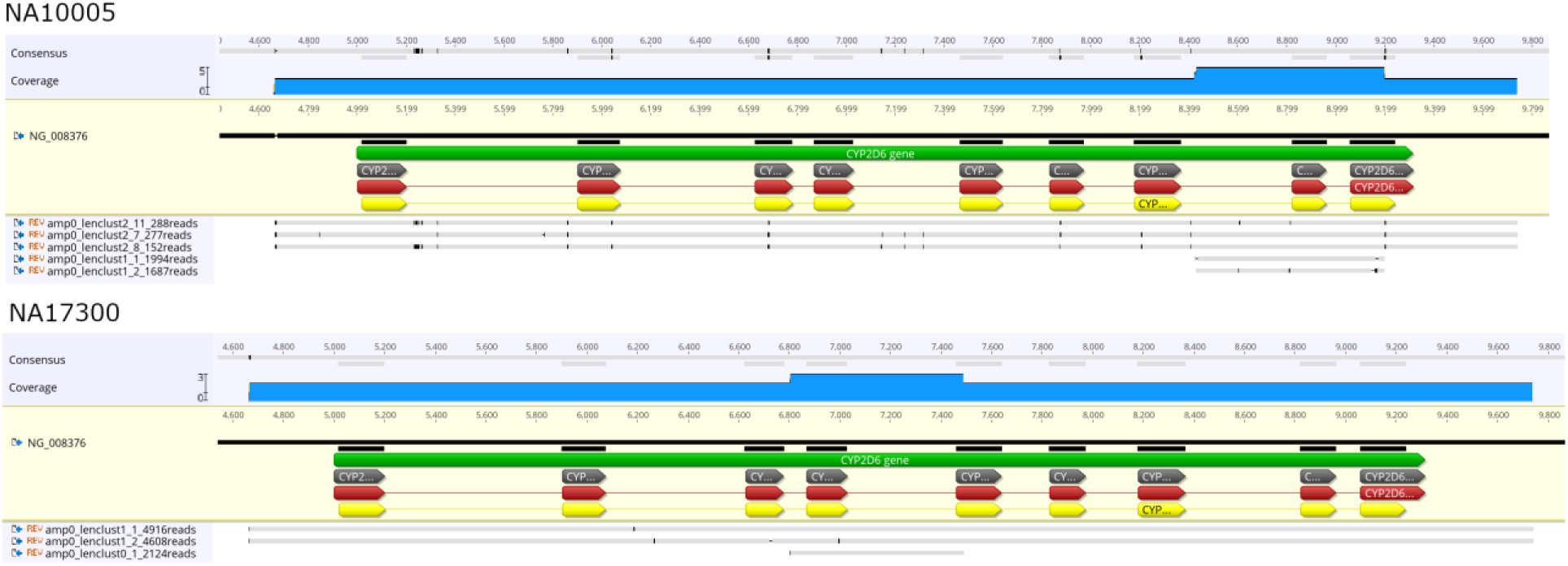
Alignment of long and short consensus sequences to *CYP2D6*. Screenshots from Geneious Prime. For NA10005 (top) and NA17300 (bottom), short consensus sequences were obtained from the respective length-clusters of short reads. The sequences of these short consensus sequences match at least one of the long consensus sequences in each sample, with the exception of mismatches at the beginning and end of the sequences.

**Supplementary Figure S3.**
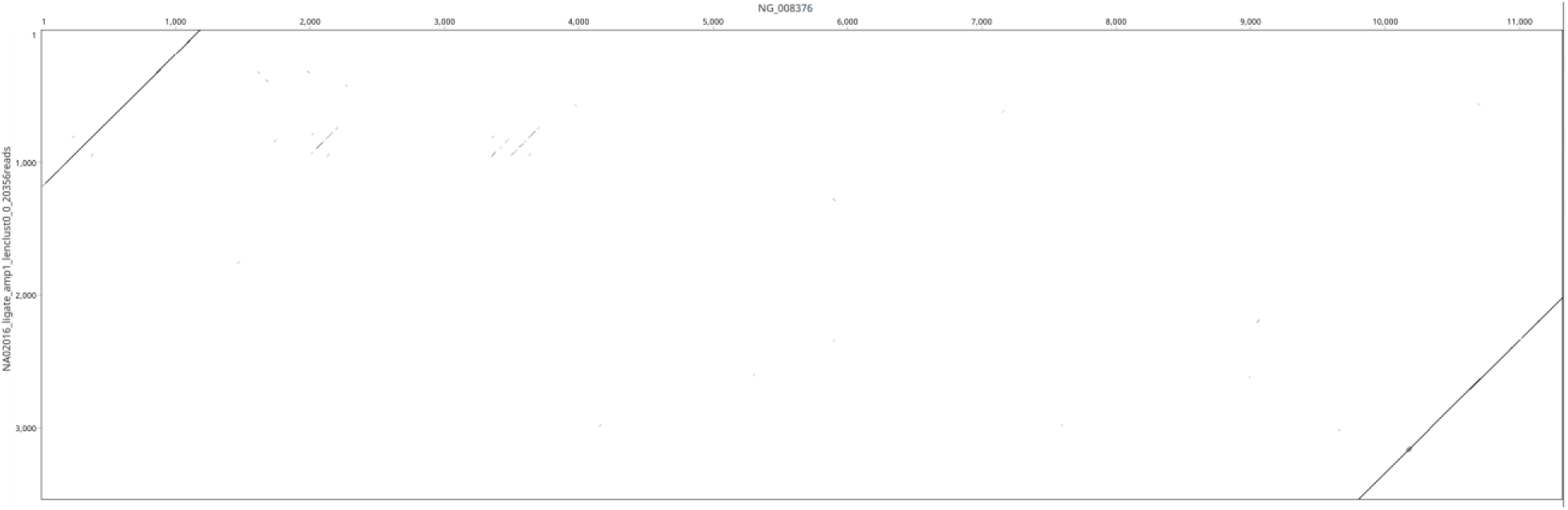
Dot plot of duplication-detecting consensus sequence and *CYP2D6*. Screenshot from Geneious Prime. The consensus sequence from NA02016_ligate from the 3.5 kb duplication-detecting amplicon (y-axis) is compared to the *CYP2D6* reference sequence (x-axis), showing the consensus sequence supporting the region upstream and downstream of *CYP2D6* as would be expected in the event of an allele duplication.

**Supplementary Figure S4.**
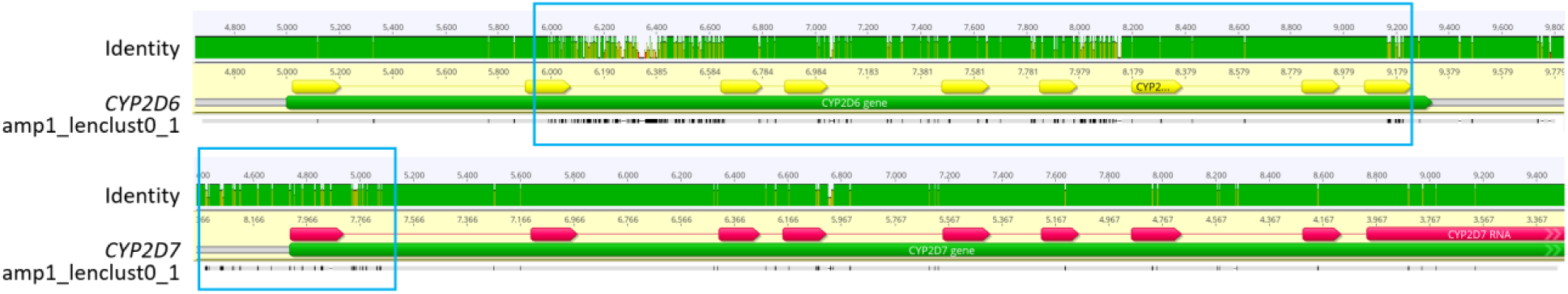
Suspected *CYP2D6-CYP2D7* hybrid allele sequence aligned to *CYP2D6* and *CYP2D7*. Screenshots from Geneious Prime. The *CYP2D7*-locus-derived consensus sequence from NA17203_ligate is aligned to the *CYP2D6* reference (top) and *CYP2D7* reference (bottom), with the sequence identity as the green histogram above the sequences. Regions of lower sequence identity to either reference are boxed in light blue. This consensus sequence appears to best support a *CYP2D6*-*CYP2D7* hybrid allele, with exon 1 from *CYP2D6*, and the remainder of the sequence from *CYP2D7*.

**Supplementary Figure S5.**
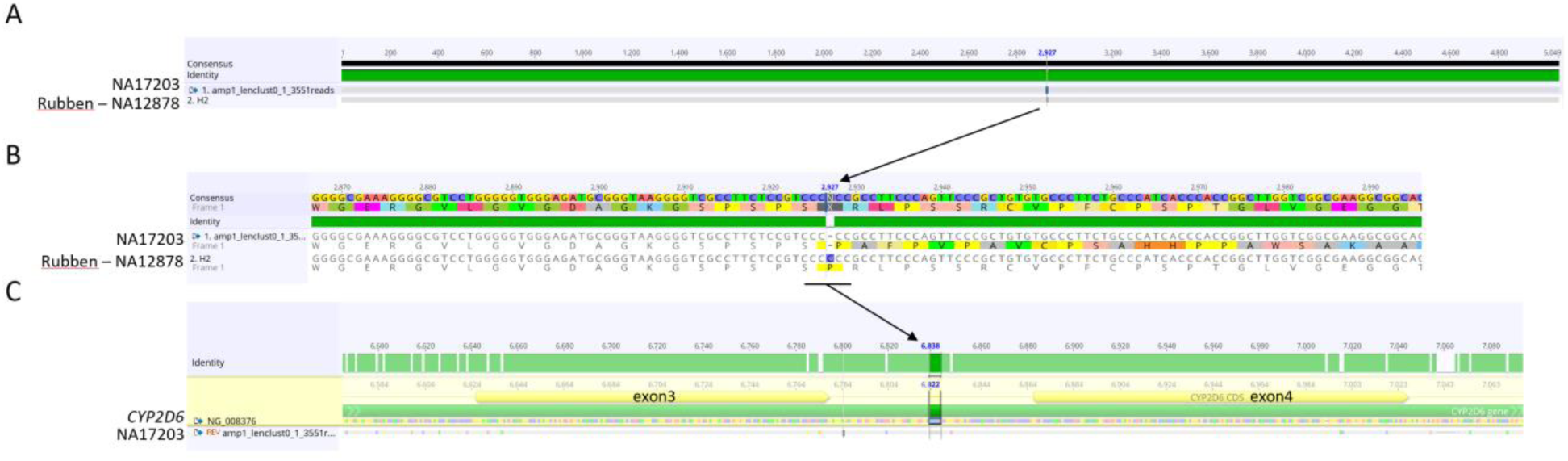
Alignment of NA17203_ligate *CYP2D6*-*CYP2D7* hybrid allele sequence to *\*68* hybrid allele sequence. Screenshots from Geneious Prime. (A) Our NA17203_ligate *\*68* allele sequence shows a perfect match to the *\*68* sequence identified from NA12878 identified by Rubben et al, (B) with the exception of a C deletion observed in our sequence within a homopolymer stretch. (C) When our *\*68* hybrid allele sequence is aligned to the *CYP2D6* reference sequence, this deletion falls between exons 3 and 4, and thus would not be expected to change the function of the resulting protein.

